# A novel missense mutation in the proprotein convertase gene *furinb* causes hepatic cystogenesis during liver development in zebrafish

**DOI:** 10.1101/2022.02.24.481764

**Authors:** Jillian L. Ellis, Kimberley J. Evason, Changwen Zhang, Makenzie N. Fourman, Jiandong Liu, Nikolay Ninov, Marion Delous, Benoit Vanhollebeke, Ian Fiddes, Jessica P. Otis, Yariv Houvras, Steven A. Farber, Xiaolei Xu, Xueying Lin, Didier Y.R. Stainier, Chunyue Yin

## Abstract

Hepatic cysts are fluid-filled lesions in the liver that are estimated to occur in 5% of the population. They may cause hepatomegaly and abdominal pain. Progression to secondary fibrosis, cirrhosis, or cholangiocarcinoma can lead to morbidity and mortality. Previous studies of patients and rodent models have associated hepatic cyst formation with increased proliferation and fluid secretion in cholangiocytes, which are partially due to impaired primary cilia. Congenital hepatic cysts are thought to originate from faulty bile duct development, but the underlying mechanisms are not fully understood. In a forward genetic screen, we identified a zebrafish mutant that develops hepatic cysts during larval stages. Cyst formation in these mutants is not due to changes in biliary cell proliferation, bile secretion, or impairment of primary cilia. Instead, time-lapse live imaging data showed that the mutant biliary cells failed to form interconnecting bile ducts because of defects in motility and protrusive activity. Accordingly, immunostaining revealed an excessive and disorganized actin and microtubule cytoskeleton in the mutant biliary cells. By whole-genome sequencing, we determined that the cystic phenotype in the mutant was caused by a missense mutation in the *furinb* gene which encodes a proprotein convertase. The mutation alters Furinb localization and causes endoplasmic reticulum (ER) stress. The cystic phenotype could be suppressed by treatment with the ER stress inhibitor 4-phenylbutyric acid and exacerbated by treatment with the ER stress inducer tunicamycin. The mutant livers also exhibited increased mTOR signaling and treatment with the mTOR inhibitor rapamycin partially blocked cyst formation by reducing ER stress. Our study has established a novel vertebrate model for studying hepatic cystogenesis and illustrated the role of ER stress in the disease pathogenesis.

## Introduction

Cystic hepatic lesions, which are characterized by progressive formation of multiple fluid-filled cysts throughout the liver, affect up to 15-18% of the population in the United States (Rawla et al., 2019). Although most patients remain clinically asymptomatic through the years, in some patients hepatic cysts may result in a massive enlargement of the liver (Santos-Laso et al., 2020). Compression of neighboring organs leads to abdominal pain and discomfort. Cyst hemorrhage, infection or rupture, as well as progression to secondary fibrosis, cirrhosis, or cholangiocarcinoma can also cause significant morbidity and mortality. Traditional treatment for symptomatic hepatic cysts includes physical removal or emptying of cysts (Drenth et al., 2010). Somatostatin analogues have been used to treat patients with severe disease but the treatment is chronic, costly, and the benefits are modest (Larusso et al., 2016). Liver transplantation remains the only definitive treatment (Santos-Laso et al., 2020).

Characterization of hepatic cysts in patients and rodent models has revealed diverse cellular defects in cholangiocytes that contribute to cyst formation, including hyperproliferation (Banales et al., 2009), enhanced fluid secretion (Banales et al., 2008), increased autophagy (Masyuk et al., 2018), changes in matrix-cell interactions caused by metalloprotease hyperactivation (Urribarri et al., 2014), and disruption in cell adhesion and polarity (Waanders et al., 2008). Meanwhile, congenital hepatic cysts that initiate in the fetal liver may result from faulty biliary development independent of cholangiocyte hyperproliferation (Beaudry et al., 2015; Raynaud et al., 2011). Cyst formation could be due to defects in hepatoblast differentiation, disturbed maturation of the ductal plate, or abnormal expansion of bile ducts (Beaudry et al., 2015; Benhamouche-Trouillet et al., 2018). Dysregulation of the signaling molecules governing biliary development can cause hepatic cystogenesis. For instance, disruption of the transcription factors HNF6 and HNF1β impairs cholangiocyte differentiation and bile duct maturation, respectively, and results in hepatic cysts (Raynaud et al., 2011).

Cholangiocytes possess primary cilia that serve as mechano-, osmo-, and chemosensors (Mansini et al., 2018). Malformation and dysfunction of primary cilia cause cholangiocyte hyperproliferation and alter their fluid secretion and bile absorption, all of which contribute to hepatic cyst formation and growth. Genetic studies have linked mutations in ciliary genes to inherited polycystic liver disease (PLD). Mutations in *PKD1* and *PKD2*, which encode the ciliary-associated proteins Polycystin-1 (PC1) and PC2, respectively, cause autosomal dominant polycystic kidney disease (ADPKD) that often manifests with liver cysts (Hughes et al., 1995; Mochizuki et al., 1996). Mutations in *PKHD1*, which encodes a primary cilium protein Fibrocystin, are associated with autosomal recessive polycystic kidney disease (ARPKD) (Nagasawa et al., 2002; Onuchic et al., 2002; Ward et al., 2002). Isolated polycystic liver diseases have been connected to mutations in genes including Protein Kinase C Substrate 80K-H (*PRKCSH*), *SEC63*, LDL receptor Related Protein 5 (*LRP5*), *SEC61B*, α-1,3-Glucosyltransferases *ALG8*, and *ALG9* (Besse et al., 2019; Besse et al., 2017; Cnossen et al., 2014; Davila et al., 2004; Li et al., 2003a). Mutations in Glucosidase II α Subunit (*GANAB*) cause rare cases of PLD with and without renal cysts (Porath et al., 2016; van de Laarschot et al., 2020). With the exception of *LRP5*, all six genes identified in isolated PLD encode proteins that are located in the endoplasmic reticulum (ER). It has been shown that mutations in *PRKCSH*, *SEC63*, *ALG8*, *SEC61B*, and *GANAB* affect the posttranslational modulation of PC1 function, raising the possibility that impairment of PC1 activity and primary cilia could be the common mechanism for PCLD (Besse et al., 2017). It is noteworthy that mutations in the known disease genes only account for 50% of clinical cases of PLD (Fabris et al., 2019; Lee-Law et al., 2019). Additional genes and mechanisms can be associated with hepatic cyst formation.

Zebrafish have become a popular model organism for studying biliary development and disease (Pham and Yin, 2019). Knocking down PCLD genes *sec63*, *prkcsh,* and *pkd1a* in zebrafish using morpholinos causes ultrastructural features of hepatic cyst formation (Tietz Bogert et al., 2013). *sec63* mutant zebrafish exhibit excessive ER stress in the liver (Monk et al., 2013). However, no genetic mutant with hepatic cysts has been reported in zebrafish.

In a forward genetic screen, we identified a novel zebrafish mutant that developed hepatic cysts at larval stages. Phenotypic analyses showed that hepatic cystogenesis in *s741* mutants was not driven by cholangiocyte hyperproliferation or defects in the primary cilium, but rather by failure to make interconnecting bile ducts during liver development. We determined that a missense mutation in the *furinb* gene, which encodes a proprotein convertase, is responsible for the biliary phenotypes. The mutation leads to mislocalization of Furinb and ER stress. Treatment with the ER stress inhibitor or chemical inhibitors of mTOR signaling partially suppressed cyst formation in the mutants.

## Results

### A forward genetic screen identifies a zebrafish mutant with hepatic nodules

To identify novel regulators of intrahepatic bile duct development, we carried out an N-ethyl-N-nitrosourea (ENU) chemical mutagenesis screen using the *Tg(EPV.Tp1- Mmu.Hbb:EGFP)^um14^/Tg(Tp1:GFP)* transgenic line that expresses GFP under the control of a Notch-responsive element. This line marks intrahepatic bile ducts and pancreatic ducts in zebrafish (Lorent et al., 2010; Parsons et al., 2009). Out of 100 F2 families screened, we isolated two recessive mutations, with one causing ductopenia and the other forming biliary nodules. *s741* mutants exhibited pericardial edema and degenerative pharyngeal arches (Fig. 1A), accompanied by 100% lethality by 1 week of age. In the liver, whereas the *Tg(Tp1:*GFP)+ cells in wild-type (WT) larvae formed a rudimental intrahepatic biliary ductal network by 96 hours post fertilization (hpf) (Fig. 1B), the *s741* mutant cells clustered in nodules and failed to form ducts (Fig. 1C,J). We performed immunostaining using the Annexin A4/Anxa4 antibody that labels intrahepatic biliary cells in zebrafish (Crosnier et al., 2005; Zhang et al., 2014). In both WT and mutant larvae, a majority of *Tg(Tp1:*GFP)+ cells were co-labeled by Anxa4, confirming that they were indeed biliary cells (Fig. 1D-I). As revealed by the nuclear labeling of Prox1, a transcription factor that is expressed in both hepatocytes and biliary cells (Chung and Stainier, 2008), the nodules in the mutants consisted of two to five biliary cells (Fig. 1I). Hematoxylin and eosin (H&E) staining showed that compared with WT siblings, the mutant livers displayed a disorganized architecture with variation in hepatocyte size and loss of the orderly arrangement of regularly-spaced hepatocytes (Fig. 1K-M). We observed scattered round to ovoid cyst-like spaces in the mutant liver at 120 hpf. Some of these spaces appeared to be entirely surrounded by hepatocytes (Fig. 1L, arrows), while others were lined by endothelium and/or contain red blood cells (Fig. 1M, arrow).

**Fig. 1.**
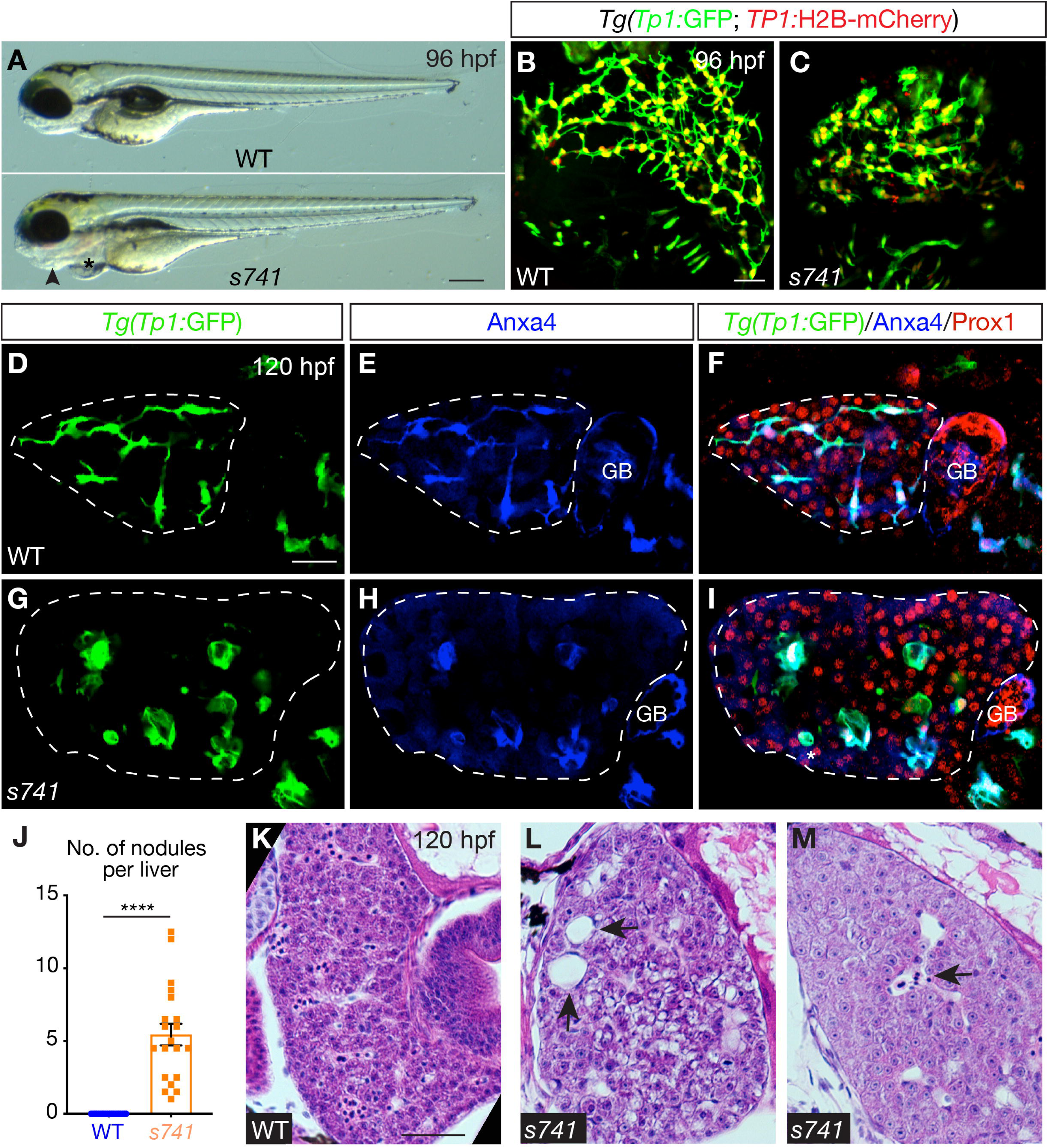
*s741* mutants develop hepatic nodules composed of intrahepatic biliary cells. (A) Live WT (top) and *s741* mutant (bottom) larvae at 96 hours post fertilization (hpf). Lateral views, anterior is to the left. Arrowhead points to the degenerating pharyngeal arches and asterisk marks the pericardial edema. (B-C) Confocal three-dimensional (3D) projections of WT and *s741* mutant larvae expressing both *Tg(Tp1:GFP)* (green) and *Tg(Tp1:H2B-mCherry)* (red) transgenes in the intrahepatic biliary cells. (D-I) Confocal single plane images of the livers in WT and *s741* mutant larvae. (D,G) *Tg(Tp1:GFP)* transgene expression marks the intrahepatic biliary cells. (E,H) Annexin A4 (Anxa4) antibody stains the biliary cells. (F,I) Merged images. Prox1 antibody labels the nuclei of hepatocytes and biliary cells. GB, gallbladder. (B-I) Ventral views, anterior is to the top. (J) Numbers of *Tg(Tp1:*GFP)+ nodules (mean±s.e.m.) in the liver of WT (blue) and *s741* mutant (orange) larvae at 96 hpf. A nodule was defined as a cluster of two or more *Tg(Tp1:*GFP)+ cells that maintained one or no interconnecting ducts with other biliary cells. Each dot represents individual liver. Statistical significance was calculated by two- tailed student’s *t*-test: ****, p<0.0001. (K-M) H&E staining of WT (K) and mutant (L,M) larval livers. Arrows in (L) point to cystic spaces that are entirely surrounded by epithelial cells (hepatyocytes and/or cholangiocytes). Arrow in (M) marks a cystic space that is lined by endothelium and contains red blood cells. Scale bars: (A) 70 μm; (B-I) 30 μm; (K-M) 50 μm.

### The intrahepatic biliary cells in *s741* mutants form cysts that retained bile fluids

To examine the bile flow in *s741* mutants, we administered fluorescent lipid analog BODIPY-FL C5:0 from 120 hpf to 128 hpf (Fig. 2A-D,G-H). BODIPY C5:0 is absorbed by the liver and secreted along with bile salts by the hepatocytes, thus allowing real-time tracking of bile secretion and flow (Carten et al., 2011). In WT, BODIPY fluorescence was detected within the intrahepatic bile ducts marked by *Tg(Tp1:*ras-mCherry) expression (Fig. 2A,C). In contrast, BODIPY fluorescence was retained within the nodules lined by the biliary cells in the mutant livers (Fig. 2B,D), indicating that these nodules are fluid-filled cysts. In WT, bile is transported through the network of intrahepatic bile ducts and drained into the gallbladder (Otis and Farber, 2013), hence, strong BODIPY fluorescence was seen in the gallbladder (Fig. 2G). While the gallbladder was formed in *s741* mutants (Fig. 2F), it was not filled with BODIPY (Fig. 2H). This is likely because bile fluids were retained in the hepatic cysts in the mutants. Patients with hepatic cysts commonly develop pancreatic and renal cysts (Bosniak and Ambos, 1975; Gabow, 1993). We found cysts in the pancreas of *s741* mutants, but not in the pronephric ducts (Supplemental Fig. S1).

**Fig. 2.**
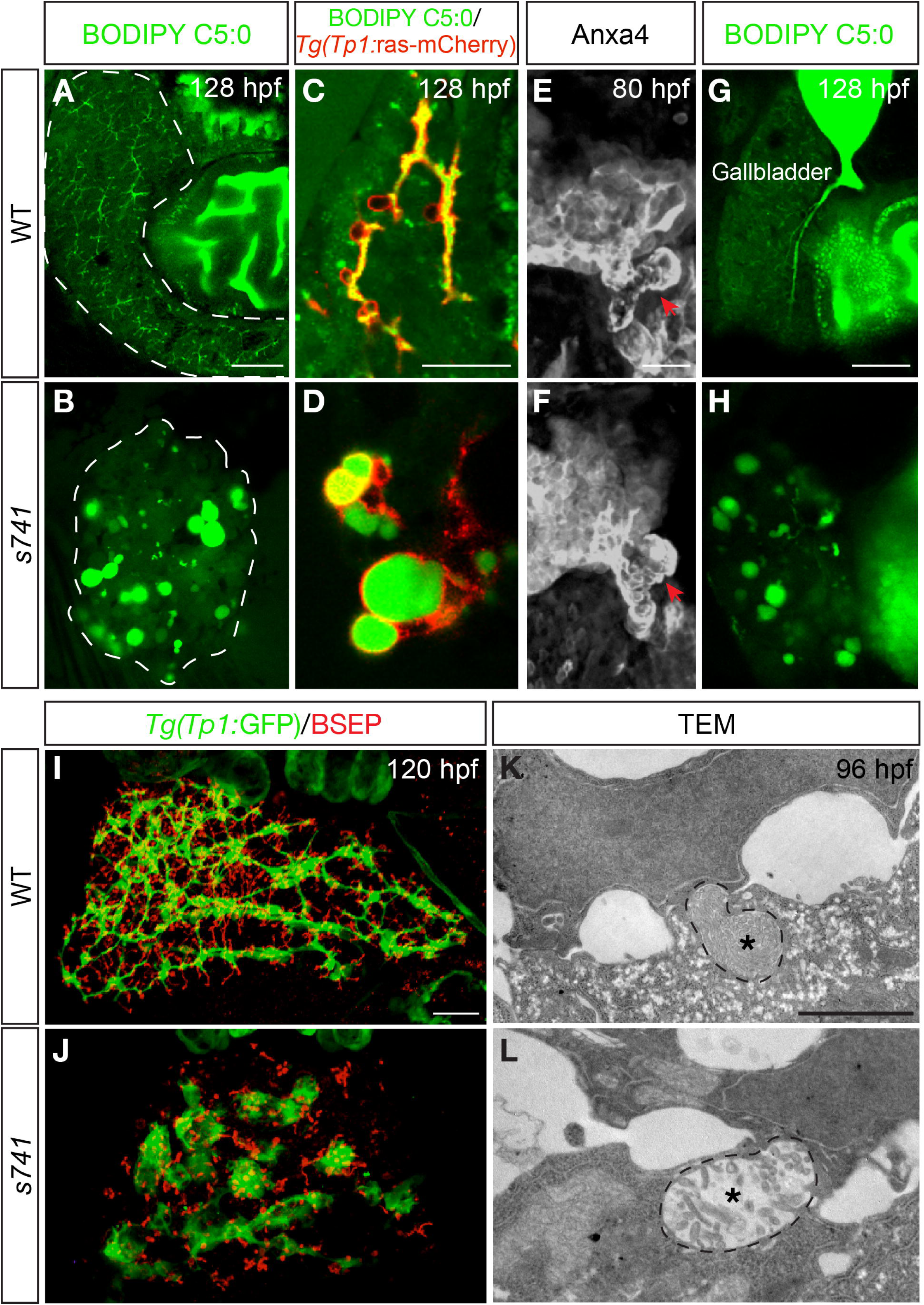
*s741* mutants form fluid-filled cysts in the liver. (A-B) Fluorescent micrographs of live WT (A) and *s741* mutant (B) larvae at 128 hpf after BODIPY feeding showing that the mutant liver retained BODIPY in the nodules. (C-D) Fluorescent micrograph images under higher magnification. These larvae also expressed *Tg(Tp1:*ras-mCherry) (red) that marked the biliary cells. 10 fish of each genotype were examined. (A-D) Lateral views taken from the left side of the fish, anterior is to the left. (E-F) Confocal 3D projections of the hepatopancreatic ductal system in WT and mutant labeled by Anxa4 antibody staining. Red arrows point to the gallbladder. Ventral views, anterior is to the top. (G-H) Fluorescent micrographs showing lack of BODIPY filling in the gallbladder of *s741* mutants. Lateral views taken from the right side of the fish, anterior is to the left. (I-J) Confocal 3D projections showing the bile ducts marked by *Tg(Tp1:*GFP) expression (green) and bile canaliculi marked by BSEP antibody staining. Ventral views, anterior is to the top. (K-L) TEM images of the bile canaliculi (dashed line) in WT (K) and *s741* mutant (L) at 96 hpf. Asterisks refer to actin microvilli within the bile canaliculi. Scale bars: (A-B, G-H) 50 μm; (C-F, I-J) 30 μm; (K-L) 2 μm.

In zebrafish, hepatocytes excrete bile salts into the bile ducts through bile canaliculi on the apical membrane (Lorent et al., 2004; Yao et al., 2012). At 120 hpf, the bile canaliculi in *s741* mutants appeared to be shorter and dilated compared to WT as revealed by the expression of bile canalicular transporter BSEP (Gerloff et al., 1998) (Fig. 2I,J). We performed transmission electron microscopy on the WT and mutant livers at 96 hpf, prior to the initiation of bile excretion in zebrafish (Otis and Farber, 2013). While the WT bile canaliculi contained tightly packed actin microvilli (Fig. 2K, asterisk), the mutant bile canaliculi were dilated and the microvilli were reduced in density and disorganized (Fig. 2L, asterisk). These observations imply that bile canalicular development was impaired in *s741* mutants independent of bile flow. It is consistent with the notion that biliary and bile canalicular development is highly coordinated (Lorent et al., 2010).

#### A missense mutation in furinb is responsible for the mutant liver phenotype

To isolate the molecular lesion responsible for the *s741* mutant phenotype, we scanned the genome for linkage to the phenotype using bulk segregant analyses and mapped the mutation to linkage group 25. We then performed whole-genome sequencing analyses. Log likelihood analysis revealed a 1.4-Mb interval, and none of the SNPs identified within the interval resulted in the gain of a new stop codon. Homozygosity scoring analysis showed two peaks within this interval (Fig. 3A) (Leshchiner et al., 2012). The first peak was located in an area where no sequence reads were returned for the mutant pool, suggestive of a deletion. The deletion covered the first 15 exons of the transcript ENSDART00000112246 that corresponds to the gene *mucin2.2/muc2.2*. The second peak was located near the transcript of the gene *furinb* and contained three non-synonymous SNPs. We generated cDNAs from 50 mutants and 50 phenotypically WT siblings, then amplified and sequenced the open reading frames of *muc2.2* and *furinb*. All the mutant embryos were homozygous for the deletion in *muc2.2* and one of the three non-synonymous SNPs of *furinb*, whereas the unaffected siblings were WT or heterozygous for these mutations (Fig. 3B and data not shown). We also sequenced the open reading frames of an additional 18 genes within the interval. There was no correlation between the presence of the nonsynonymous or splice variants in these genes and the mutant phenotypes (data not shown).

**Fig. 3.**
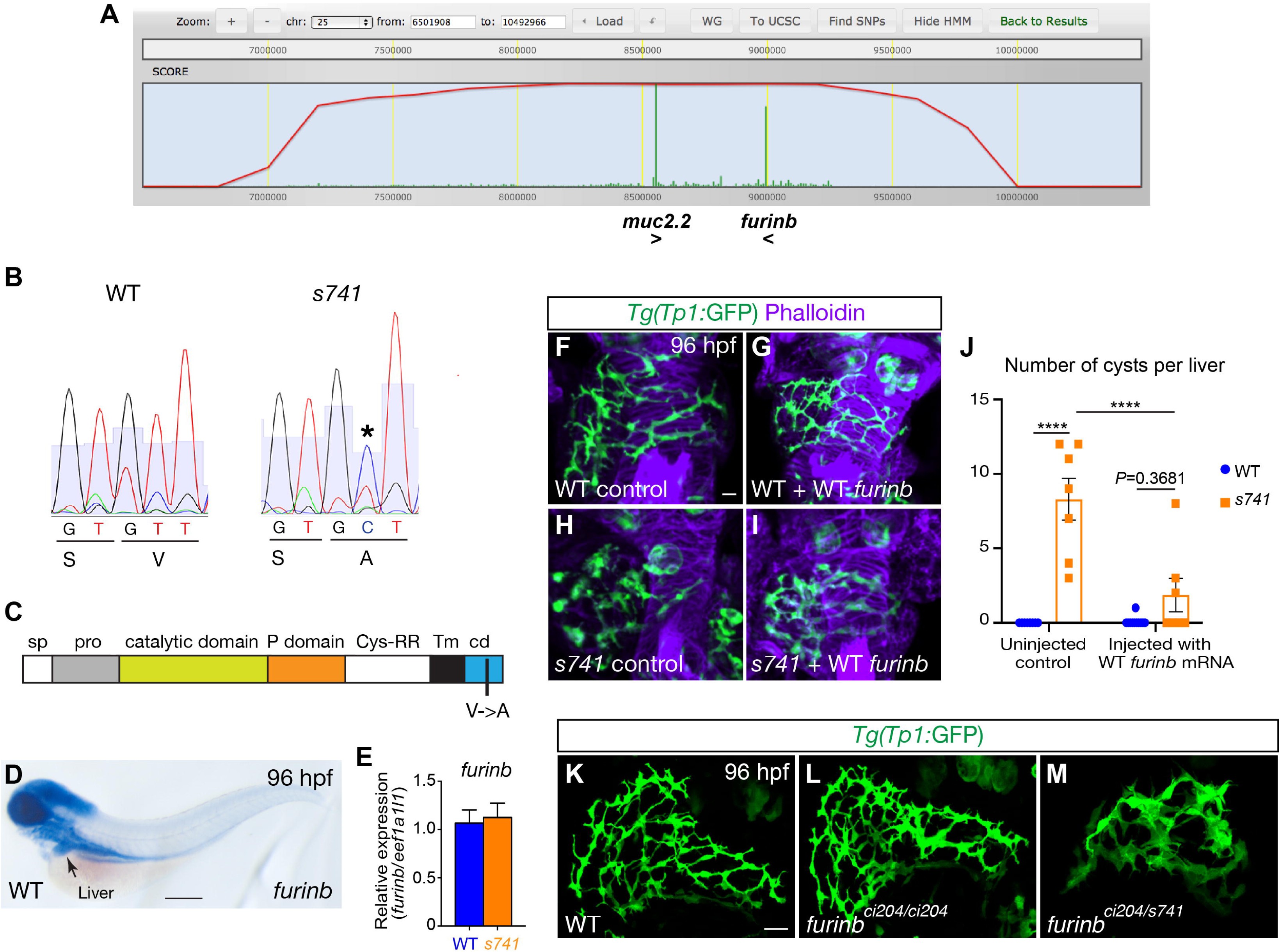
A missense mutation in the *furinb* gene is responsible for the cystic phenotype in *s741* mutants. (A) Genome view of linkage analysis using SNPtrack. Log likelihood analysis (red line) returned a ∼1.4-Mb interval on linkage group 25 for the presumptive mutation. The homozygosity score (green) suggested two candidate mutations in the *muc2.2* and *furinb* genes. (B) Sequencing of cDNA from *s741* mutants and WT siblings. Mutants bear a T-to-C mutation (indicated by *) near the C terminus, leading to a V-to-A amino acid change. (C) Domain diagram of Furinb protein. The V-to-A change occurs within the cytoplasmic domain (cd). Sp, signal peptide; pro, pro-domain; Cys-RR, cysteine-rich region; Tm, transmembrane domain. (D) Whole-mount *in situ* hybridization detected *furinb* transcript in the head and digestive organs in WT at 96 hpf. Lateral view, anterior is to the left. (E) qPCR analysis comparing *furinb* transcripts in the WT and *s741* mutant larval livers at 96 hpf. Triplicates were performed. The results are represented as relative expression levels normalized to the housekeeping gene *eef1a1l1* (mean±s.e.m.). Statistical significance was calculated by two-tailed student’s *t*-test: p=0.7877. (F-I) Confocal 3D projections showing uninjected control WT and *s741* mutant larvae (F,H) and larvae injected with WT *furinb* mRNA at the one-cell stage (G,I). Phalloidin (purple) stained for F-actin and *Tg(Tp1:*GFP) expression (green) labeled the intrahepatic biliary cells. (J) Numbers of cysts per liver (mean±s.e.m.) in control and WT *furinb* mRNA- injected larvae. Each dot represents an individual liver. Statistical significance was calculated by one-way ANOVA and Tukey’s post-hoc test: ****, p<0.0001. (K-M) Confocal 3D projections showing the livers in WT (K), *furinb^ci204/ci204^* CRISPR mutant (L), and *furinb^ci204/s741^* compound heterozygote (M). The intrahepatic bile ducts were marked by *Tg(Tp1:*GFP) expression. At least five fish were examined for each genotype. (F-I, K-M) Ventral views, anterior is to the top. Scale bars, (D) 70 μm; (F-I, K-M) 20 μm.

To determine which mutation was responsible for the cystic phenotype in *s741* mutants, we first examined the expression of *muc2.2* and *furinb* transcripts. *muc2.2* encodes a member of the Mucin protein family that are the main gel-forming constituent of mucus (Voynow and Rubin, 2009). Consistent with a previous report (Jevtov et al., 2014), we detected *muc2.2* transcript in the testes of adult male zebrafish by reverse transcription PCR. However, there was no *muc2.2* expression in the livers of WT or *s741* mutant larvae at 96 hpf (Supplemental Fig. S2A). We also generated a *muc2.2* mutant allele that resembled the deletion mutation found in *s741* mutants by CRISPR/Cas9 genome editing (Supplemental Fig. S2B). These mutants did not have obvious bile duct phenotypes at 96 hpf (Supplemental Fig. 2C-D). Therefore, the *muc2.2* mutation unlikely accounts for the *s741* mutant phenotypes.

*furinb* is one of the zebrafish orthologs of the mammalian *Furin* gene that encodes an endoprotease belonging to a family of proprotein convertases (Walker et al., 2006). The missense mutation in *furinb* found in *s741* mutants causes a valine to alanine amino acid change at the C-terminus of the Furinb protein (p.V822A) (Fig. 3B-C). Whole mount *in situ* hybridization showed that *furinb* transcript is expressed in the central nervous system, eye, heart, jaw, and digestive organs during development (Fig. 3D, Anderson et al., 2013). Quantitative real-time PCR (qPCR) showed that the *s741* mutant livers had comparable expression of *furinb* as WT at 96 hpf (Fig. 3E). Injection of WT *furinb* mRNA significantly reduced the number of hepatic cysts in *s741* mutants, confirming that the missense mutation in the *furinb* gene is responsible for the cystic phenotype (Fig. 3F-J). To test if Furinb is required for bile duct development, we generated *furinb* null mutants by CRISPR/Cas9 genome editing. The *furinb^ci204^* mutant harbors a 17bp deletion in the second exon leading to a premature stop codon (Supplemental Fig. S3A). The resulting protein is predicted to truncate at the beginning of the propeptide domain (Supplemental Fig. S3B). *furinb^ci204^* mutants did not show bile duct phenotypes at 96 hpf (Fig. 3L). We crossed *s741* heterozygotes with *furinb^ci204^* heterozygotes and examined the intrahepatic bile ducts in the progeny. The genotypes of the progeny were determined by sequencing. Neither *s741* heterozygous nor *furinb^ci204^* heterozygous fish exhibited evident abnormality in bile duct morphology (data not shown), but the compound heterozygotes showed clustering of the biliary cells (Fig. 3M). Unlike *s741* mutants, the compound heterozygotes did not form hepatic cysts. Thus the *furinb* missense mutation found in *s741* mutants is likely a neomorphic allele.

#### The furinb missense mutation impairs bile duct development in a cell-autonomous manner

To determine if the cystic phenotype seen in *s741* mutants was due to abnormal Furinb function within the biliary cells or their environment, we performed genetic mosaic analyses (Fig. 4). We injected the WT donor embryos with *casanova/sox32* mRNA at the one-cell stage to render the cells towards an endodermal fate (Kikuchi et al., 2001; Sakaguchi et al., 2001). We transplanted cells from the dorsolateral margin of the WT donor embryos into the blastodermal margin of the *s741* mutant host embryos at 40% epiboly, and harvested the host larvae at 100 hpf (Fig. 4A). The WT donor cells formed bile ducts with normal appearance in the mutant host liver (Fig. 4B). In the converse experiments, we transplanted the *s741* mutant cells into the WT host embryos. The mutant cells still formed cysts in the WT liver (Fig. 4C). We concluded that the mutant Furinb protein acts within the biliary cells to drive cyst formation.

**Fig. 4.**
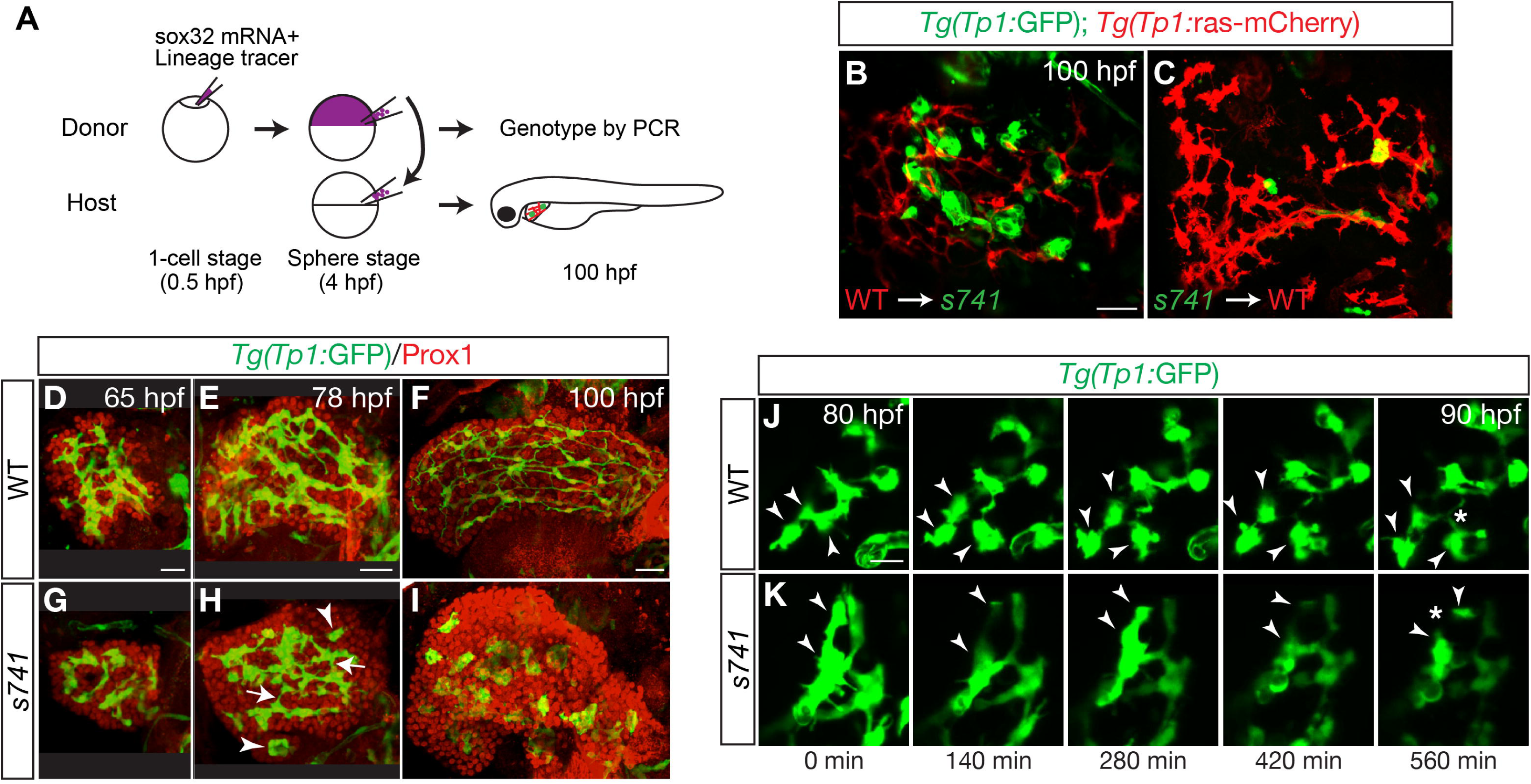
The mutant Furinb protein acts within the biliary cells to disrupt bile duct morphogenesis. (A) Schematic diagram of the genetic mosaic analysis for determining the cell autonomy of the mutant Furinb function. (B-C) Representative results of the genetic mosaic analysis. (B) shows WT donor cells transplanted into the *s741* mutant host liver (6 host fish examined). (C) shows *s741* mutant donor cells transplanted into the WT host liver (10 host fish examined). In both B and C, the WT biliary cells expressed *Tg(Tp1:*ras-mCherry) (red) and the *s741* mutant biliary cells expressed *Tg(Tp1:*GFP) (green). (D-I) Time-course analysis of bile duct morphogenesis. *Tg(Tp1:*GFP) expression marks the intrahepatic biliary cells (green), and Prox1 antibody labels the nuclei of hepatocytes and biliary cells (red). In (H), arrows point to the interconnecting ducts between biliary cells. Arrowheads mark biliary cell clusters that did not connect to other biliary cells. (B-I) Confocal 3D projections. Ventral view, anterior is to the top. (J-K) Snapshots of time-lapse live imaging of *Tg(Tp1:*GFP)+ biliary cells in WT (J) and *s741* mutant (K) at the time points indicated. Arrowheads in (J) and (K) in each panel point to the same group of biliary cells over time. Asterisk in (J) shows the interconnecting ducts formed among the three biliary cells marked by arrowheads. Asterisk in (K) shows the loss of the pre-existing connecting duct between the two biliary cells marked by arrowheads. Lateral view, anterior is to the left. Confocal three-dimensional projections are shown. Scale bars: (B-I) 30 μm; (J,K) 20 μm.

### The biliary cells in *s741* mutants fail to undergo proper morphogenesis to make bile ducts

To identify the morphogenic events that lead to hepatic cyst formation in *s741* mutants, we performed time-course analyses by tracking *Tg(Tp1:*GFP)+ cells in WT and mutant larvae that were harvested at various time points during liver development. At 65 hpf, the *Tg(Tp1:*GFP)+ cells in WT and *s741* mutant livers displayed similar appearance (Fig. 4D,G). At 78 hpf, the *Tg(Tp1:*GFP)+ populations in both the WT and mutant livers had expanded (Fig. 4E,H). In WT, they interconnected via cellular protrusions and formed a rudimental intrahepatic biliary network (Fig. 4E). In the mutant, some cells had formed connecting ducts (Fig. 4H, arrows), whereas the others were clustered in isolated nodules (Fig. 4H, arrowheads). Between 78 and 100 hpf, the *Tg(Tp1:*GFP)+ cells in the WT liver continued to spread while remaining interconnected to construct a complex biliary ductal network (Fig. 4F). In contrast, the mutant cells formed nodules rather than interconnecting ducts (Fig. 4I).

To characterize the cellular behaviors of the biliary cells in real time, we conducted time-lapse live imaging between 80 and 90 hpf. In WT (Fig. 4J, supplemental movie 1), the *Tg(Tp1:*GFP)+ biliary cells were seen in clusters at the beginning of the time lapse. Over the next ∼10 hours, individual biliary cells migrated away from each other but at the same time sent out protrusions that interconnected to form bile ducts (Fig. 4J, arrowheads and asterisk, Lorent et al., 2010). In the mutant, the biliary cells had very little motility throughout the imaging time period (Supplemental movie 2). Instead of extending new protrusions to make bile ducts, some cells even retracted the pre-existing connections (Fig. 4K, arrowheads and asterisk), leading to nodule formation.

In mammals, excessive proliferation of cholangiocytes contributes to the initiation and progressive growth of hepatic cysts (Fabris et al., 2019). To evaluate the proliferation of biliary cells, we incubated WT and mutant larvae with the replication marker 5- ethynyl-2’-deoxyuridine (EdU) from 80 to 96 hpf and analyzed EdU incorporation in the *Tg(Tp1:*GFP)+ biliary cells (Fig. S4A). The percentage of EdU-positive biliary cells was only slightly higher in the mutant compared to WT, and the difference was not statistically significant. We also performed TUNEL assay and only detected sporadic apoptotic cells in WT and mutant livers at 96 hpf (Supplemental Fig. S4B-E). Therefore, dysregulated proliferation or apoptosis does not account for cyst formation in *s741* mutants.

### Distribution of the actin and microtubule cytoskeleton is altered in the *s741* mutant biliary cells

The defects in cell motility and protrusive activity prompted us to examine the actin and microtubule cytoskeleton in the *s741* mutant biliary cells. Phalloidin staining showed that in the WT liver, F-actin was located at the cell periphery and enriched in the bile canaliculi (Fig. 5A,A’). There seemed to be an excessive accumulation of F-actin within the *s741* mutant biliary cells compared to WT (Fig. 5B,B’). There was also an ectopic accumulation of stabilized microtubules in the mutant biliary cells as revealed by immunostaining with an anti-Acetylated tubulin antibody (Fig. 5C-D’).

**Fig. 5.**
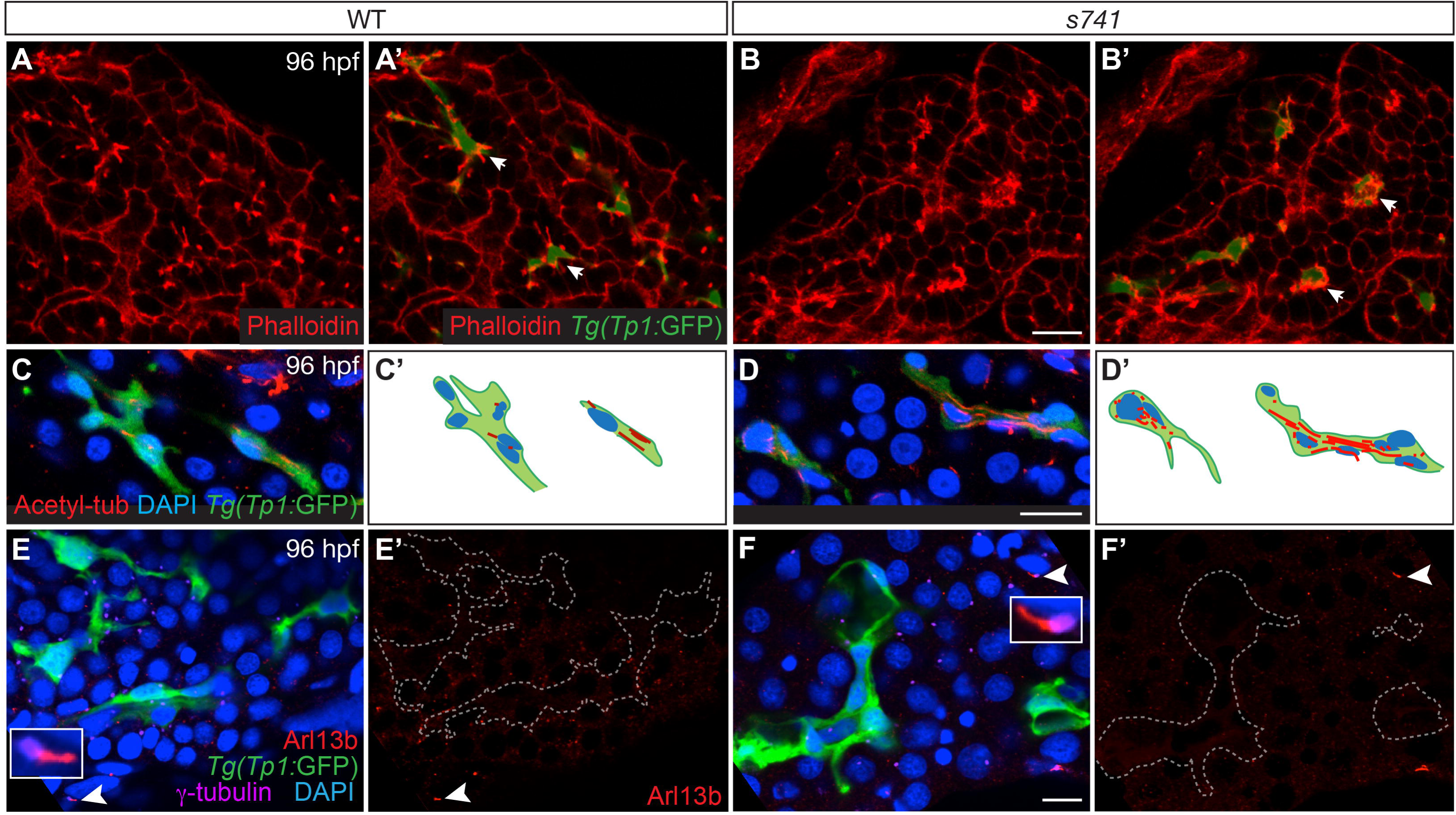
Distribution of F-actin and microtubule cytoskeleton is altered in the *s741* mutant biliary cells. (A,B) Confocal single plane images of WT and *s741* mutant livers stained with Phalloidin to label F-actin. (A’,B’) are the same samples as (A,B) with *Tg(Tp1:*GFP) expression marking the biliary cells (green). (C,D) Confocal single plane images of WT and *s741* mutant livers stained with acetylated-tubulin/acetyl-tub antibody for stabilized microtubules (red), DAPI for nucleus (blue), and *Tg(Tp1:*GFP) expression for biliary cells (green). (C’,D’) Schematic diagrams of the biliary cells shown in (C,D). Nuclei and acetylated-tubulin staining are indicated. (E,F) Confocal single plane images of WT and *s741* mutant livers stained with Arl13b antibody for primary cilia (red), γ-tubulin for basal bodies (purple), DAPI for nucleus (blue), and *Tg(Tp1:*GFP) expression for biliary cells (green). (E’,F’) show the same samples as in (E,F) but with only Arl13b staining. The biliary cells, which are outlined by dashed lines, lacked obvious Arl13b staining, whereas the neighboring cells had distinct primary cilia that were marked by Arl13b (white arrowheads in E,E’,F,F’). Inserts in E and F show high magnification images of representative primary cilia marked by arrowheads. (A-F) Vibratome sections. Scale bar: (A,B) 20 μm; (C-F) 10 μm.

In mammals, hepatic cystogenesis has been associated with defects in the primary cilium (Lee and Somlo, 2014). We used an anti-Arl13b antibody to specifically mark the primary cilia (Duldulao et al., 2009) and an anti-γ-tubulin antibody to label the basal bodies (Lunt et al., 2009). We detected primary cilia and associated basal bodies in the mesenchymal cells in WT and mutant livers (Fig. 5E-F’, arrowheads and inserts). However, we failed to detect Arl13b staining in proximity to the basal bodies in the biliary cells in the WT or mutant animals (Fig. 5E-F’). In patients with ADPKD, inactivation of PC1 impairs primary cilia function and leads to cystogenesis in the kidney and liver (Lee and Somlo, 2014). We obtained zebrafish *pkd1a* individual mutants and *pkd1a;pkd1b* compound mutants which develop pronephric cysts at larval stages (Zhu et al., 2017). They did not form hepatic cysts (Supplemental Fig. S5). These results suggest that hepatic cystogenesis in *s741* mutants is unlikely due to the gross impairment of primary cilia.

### The *s741* missense mutation alters Furinb localization and triggers inflammation and the unfolded protein response

Given that the *furinb* missense mutation found in *s741* mutants did not alter the transcript expression, we asked if it changed the protein expression. As Furinb-specific antibody was not available, we generated transgenic constructs that utilized the Notch reporter *Tp1* to drive the expression of either WT or *s741* mutant Furinb-GFP fusion protein in the biliary cells. We injected the constructs into WT embryos at the one-cell stage and performed live imaging of their livers at 120 hpf. Whereas the WT Furinb-GFP fusion protein was located along the intrahepatic bile ducts (Fig. 6A-C), the mutant Furinb-GFP fusion protein was localized in puncta within the biliary cells (Fig. 6D-F). Moreover, the WT biliary cells expressing the mutant Furinb-GFP fusion protein tended to lose their ductular morphology and became clustered together (Fig. 6E, arrows).

**Fig. 6.**
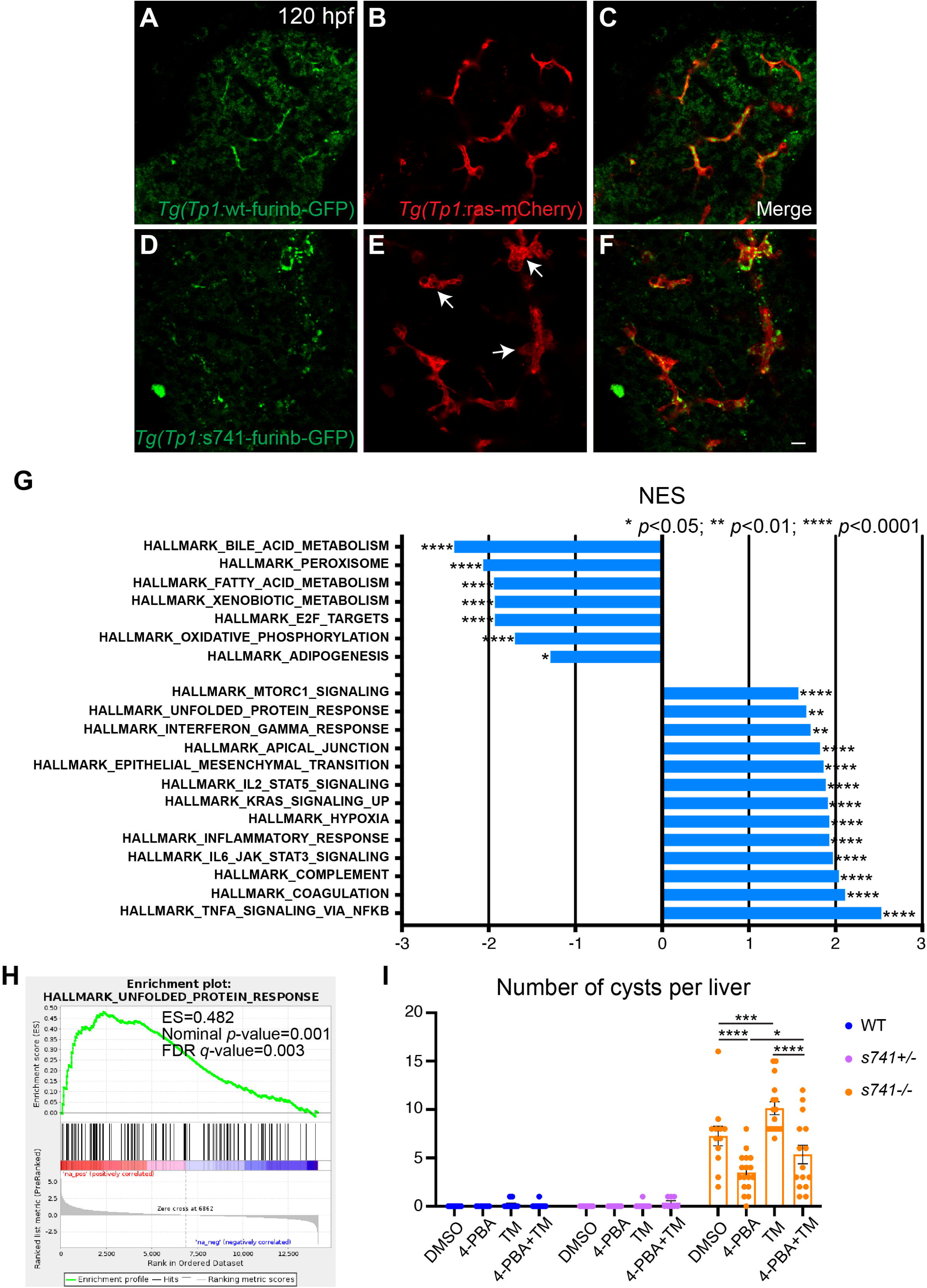
The *s741* mutation causes mislocalization of Furinb, which triggers ER stress. (A-F) Confocal single plane images showing the expression of WT and *s741* mutant Furinb- GFP fusion proteins in the biliary cells. (A,D) show only the Furinb-GFP fusion protein, (B,E) show biliary cells marked by *Tg(Tp1:*ras-mCherry) expression, and (C,F) show merged images. Arrows in E mark clusters of biliary cells that are not seen in WT fish expressing WT furinb-GFP fusion protein. Scale bar, 10 μm. (G) Gene set enrichment analysis (GSEA) identified gene sets that were significantly altered in *s741* mutant livers with nominal p-value <0.05 and false discovery rate (FDR)<0.01. Gene sets were ranked by normalized enrichment score (NES). (H) GSEA plots showing a significant enrichment of unfolded protein response (UPR) pathway in *s741* mutant livers compared to WT livers. (I) Numbers (mean±s.e.m.) of hepatic cysts as revealed by *Tg(Tp1:*GFP) expression in WT, *s741+/-* heterozygotes, and *s741-/-* homozygous mutants after treatments with 10 mM 4-4-phenylbutyric acid (4-PBA), 1 mg/mL tunicamycin (TM), or both from 72 to 120 hpf. Each dot represents individual liver. Statistical significance was calculated by one-way ANOVA and Tukey’s post-hoc test. *, p<0.05; ***, p<0.001; ****, p<0.0001.

To uncover the molecular mechanisms underlying the cystic phenotype in *s741* mutants, we performed global transcriptomic analysis on the livers of WT and *s741* mutant larvae at 100 hpf. Gene set enrichment analysis (GSEA) revealed reduction in metabolism pathways and upregulation of TNFα signaling, IL6-JAK-STAT3 signaling, and hypoxia genes in the mutant livers (Fig. 6G), indicating that the accumulation of mutant Furinb protein compromised liver function and provoked an inflammatory response, respectively. Genes involved in unfolded protein response (UPR) were upregulated in the mutant livers (Fig. 6H), which was correlated to the mislocalization of the mutant Furinb protein. To determine if increased UPR and ER stress contributed to the cystic phenotype in the mutant, we treated the mutants and their WT and heterozygous siblings with the ER stress inhibitor 4-phenylbutyric acid (4-PBA) (Ozcan et al., 2006), the ER stress inducer tunicamycin (TM) (Banerjee et al., 2011), or both chemicals from 72 to 120 hpf (Fig. 6I). None of the chemical treatments had obvious impact on the bile ducts in WT and *s741* heterozygous larvae. Meanwhile, inhibition of ER stress by 4-PBA significantly reduced the number of hepatic cysts in *s741* mutants. Exacerbated ER stress induced by TM increased the number of hepatic cysts in the mutants, and such an increase was suppressed by dual treatment with 4-PBA and TM. The results from the chemical treatment support the role of ER stress in cyst formation in *s741* mutants.

RNAseq analysis also indicated that the mutant livers had more inflammation compared to WT. Consistently, we detected a more than 5-fold increase in the number of *Tg(mpeg1:*YFP)+ macrophages in the mutant livers compared to WT at 120 hpf (Supplemental Fig. S6A,C,F) (Roca and Ramakrishnan, 2013). However, at 96 hpf when the cystic phenotype first became evident in the mutants, there was no significant difference in the numbers of macrophages between WT and mutants (Fig. S6E). Thus increased inflammatory response does not contribute to the initiation of cystogenesis in *s741* mutants. However, it may play a role in the progression of bile duct injury in these animals.

### Treatment with the mTOR inhibitor rapamycin partially rescues the cystic phenotype in *s741* mutants

To seek signaling pathways that can be targeted to suppress hepatic cystogenesis, we focused on mTOR signaling. It is upregulated in the mutant liver as revealed by RNA-seq (Fig.. 7A) and has been shown to mediate both ER stress and inflammation (Saxton and Sabatini, 2017). To test if inhibition of mTOR signaling has any effect on hepatic cystogenesis in *s741* mutants, we treated the mutants and their WT siblings with DMSO or 5 μM rapamycin from 72 to 120 hpf and examined the morphology of the intrahepatic bile ducts. Rapamycin treatment did not significantly change the total number of biliary cells in WT (*p*=0.9903) or *s741* mutants (*p*=0.6595). Meanwhile, it reduced the number of hepatic cysts in the mutants compared to the DMSO-treated control (Fig. 7B-F).

**Fig. 7.**
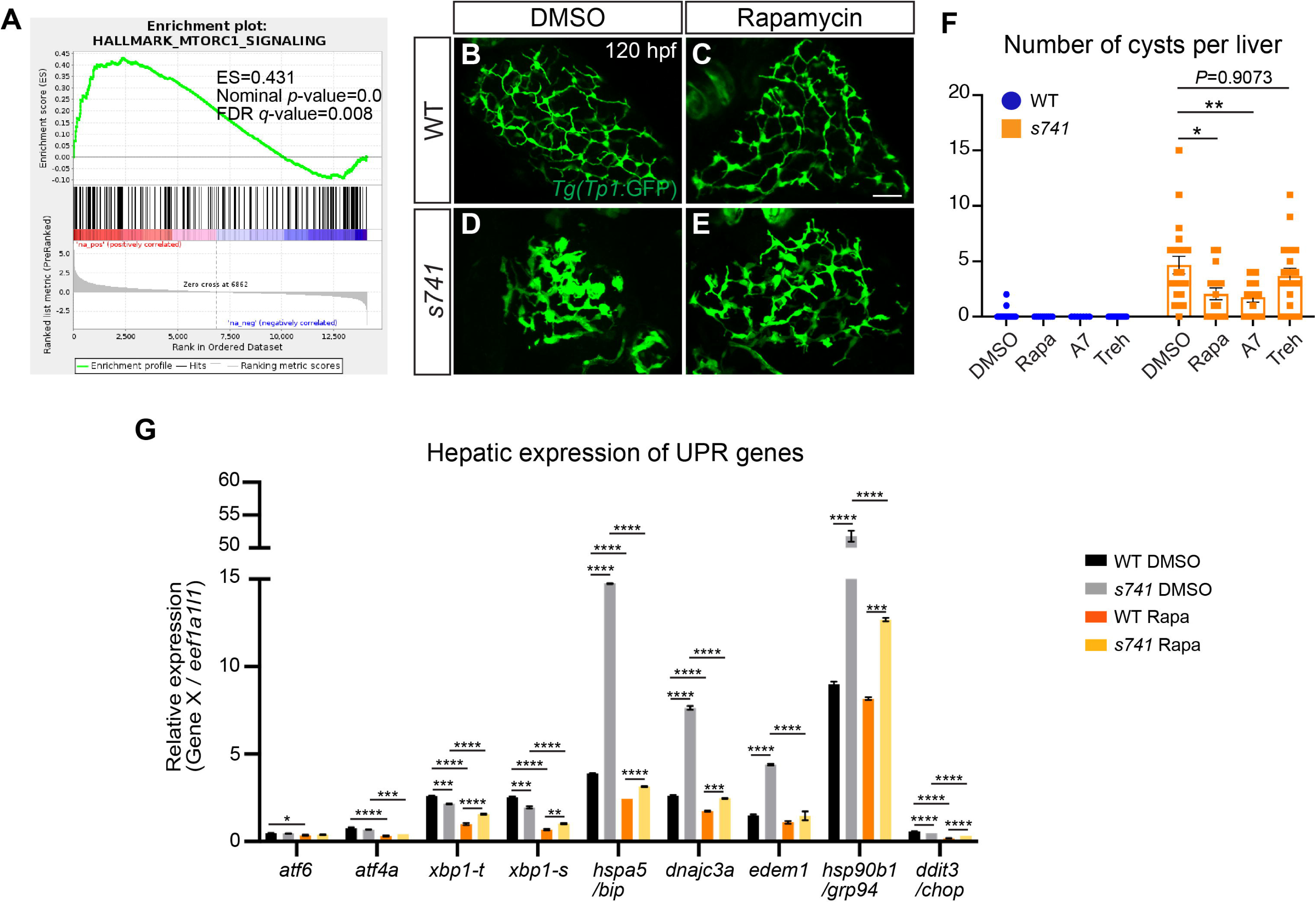
Rapamycin treatment reduces the number of hepatic cysts in *s741* mutants. (A) GSEA plot showing a significant enrichment of mTORC1 signaling in *s741* mutant livers. (B-E) Confocal 3D projections of WT and *s741* mutant livers after DMSO or 5 μM rapamycin treatment from 72 to 120 hpf. The biliary cells are marked by *Tg(Tp1:*GFP) expression. Ventral views, anterior is to the top. Scale bar, 30 μm. (F) Numbers (mean±s.e.m.) of liver cysts as revealed by *Tg(Tp1:*GFP) expression in WT and *s741* mutants after chemical treatments from 72 to 120 hpf. Each point represents an individual liver. (G) qPCR analyses showing the hepatic expression of UPR response genes in WT and *s741* mutants after DMSO or rapamycin treatment from 72 to 120 hpf. Triplicates were performed. The results are represented as relative expression normalized to the housekeeping gene *eef1a1l1* (mean±s.e.m.). Statistical significance in (F,G) was calculated by one-way ANOVA and Tukey’s post-hoc test. *, p<0.05; **, p<0.01; ***, p<0.001; ****, p<0.0001. Rapa, rapamycin; A7, A769662; Treh, trehalose.

Treatment with 10 μM A769662 (Goransson et al., 2007), which suppresses mTOR through activating its upstream negative regulator AMP-activated protein kinase (AMPK), resulted in a similar reduction in cyst numbers in the mutants (Fig. 7F). Rapamycin is a potent inducer of autophagy (Noda and Ohsumi, 1998). However, treatment with 2% trehalose, which is an mTOR-independent autophagy inducer (Mardones et al., 2016), did not reduce the number of hepatic cysts in the mutants, suggesting that rapamycin may not suppress hepatic cystogenesis through inducing autophagy. To test if rapamycin treatment acted through reducing ER stress, we performed qPCR on the control or rapamycin-treated larvae from 72 to 120 hpf. Consistent with the RNA-seq data, the hepatic expression levels of UPR effector genes, including *bip*, *dnajc3a*, *edem1*, and *grp94*, were significantly increased in *s741* mutants compared to WT (Fig. 7G). Such increases were reversed by rapamycin treatment (Fig. 7G). Rapamycin treatment also reduced the expression of pro-inflammatory genes and the number of macrophages in the *s741* mutant livers (Supplemental Fig. S6B,D,F,G). Taken together, treatment with mTOR inhibitors partially suppressed hepatic cystogenesis in *s741* mutants, which coincided with the reduction in ER stress and inflammation in the mutant livers.

## Discussion

Using forward genetics, we identified the first zebrafish genetic mutant that forms hepatic cysts during liver development. Hepatic cystogenesis in *s741* mutants is independent of cholangiocyte hyperproliferation or ciliopathies. Instead, changes in the actin and microtubule cytoskeleton perturb the protrusive behaviors of the biliary cells and prevented them from forming interconnecting bile ducts. We attribute the morphogenetic defects to the mislocalization of proprotein convertase Furinb caused by a missense mutation in the C-terminus and the subsequent induction of ER stress. Hepatic cystogenesis was partially suppressed by treatment with the ER stress inhibitor 4-PBA. Our transcriptomic analysis revealed an increase of mTORC1 signaling in *s741* mutants. Treatment with mTOR inhibitors ameliorated cyst formation in the mutants, at least in part by reducing ER stress.

Studies in human and rodent models have shown that congenital hepatic cystogenesis originates from malformation of the ductal plate during embryogenesis (Benhamouche-Trouillet et al., 2018; Raynaud et al., 2011). In fetal liver tissues from patients with ARPKD and Cpk mice that carry mutations in the *Cystin1* gene encoding cilium-associated protein, perturbed apico-basal polarity of the cholangiocytes is associated with dysmorphogenesis of the ductal plate and cyst formation (Raynaud et al., 2011). Cpk mice exhibit excessive and accelerated differentiation of hepatoblasts into cholangiocyte precursors, which may initiate hepatic cystogenesis (Benhamouche-Trouillet et al., 2018). Our study of *s741* mutants provides another example that defective bile duct development serves as the initiating cause of liver cysts during embryogenesis. The biliary cells in *s741* mutants had reduced motility and protrusive activity and formed isolated nodules rather than interconnecting bile ducts.

In humans, PLDs have been linked to defects in the structure or function of primary cilia in cholangiocytes (Masyuk et al., 2021). Our study suggests that ciliopathy may not cause hepatic cystogenesis in zebrafish during development. Immunostaining using the primary cilia marker Arl13b antibody failed to detect primary cilia in biliary cells at the larval stage. Despite of developing renal cysts as seen in patients with ADPKD (Zhu et al., 2017), *pkd1* mutant larvae did not form liver cysts. *pkd2* mutant zebrafish did not develop liver cysts either (Schottenfeld et al., 2007; Sun et al., 2004) (Zhaoxia Sun, personal communication). It is noteworthy that *pkd1* and *pkd2* mutant zebrafish are premature lethal. Whether primary cilia play a role in cholangiocyte physiology in adult zebrafish remains to be determined.

Instead of defects in primary cilia, we attribute the liver cyst formation in *s741* mutants to changes in the actin and microtubule cytoskeleton. Disorganized actin cytoskeleton has been observed in cystic *PKD1*-null kidney epithelial cells derived from patients with ADPKD (Streets et al., 2020). Both PC1 and PC2 have been shown to interact directly with actin cytoskeleton molecules and mediate actin cytoskeleton organization and directional cell migration *in vitro* (Boca et al., 2007; Gallagher et al., 2000; Li et al., 2003b; Yao et al., 2014). Similar to what was seen in *s741* mutant biliary cells, ARPKD human kidney tissue has excessive acetylated-tubulin (Berbari et al., 2013). Treatment of microtubule-specific agents colchicine, vinblastine, and Taxol inhibits renal cyst development *in vitro* independent of their antimitotic function (Woo et al., 1994). It will be interesting to investigate to what extent changes in actin and microtubules contribute to hepatic cyst formation. Our current hypothesis is that disruption of actin and microtubule cytoskeleton impairs cell-cell adhesion and protrusive activity essential for bile duct morphogenesis, leading to cystogenesis. Both actin and microtubules play important roles in protein trafficking and polarized protein localization. In the future, we will investigate if disruption of actin and microtubules alters the expression of channel and transporter proteins in biliary cells to enhance cyst formation (Avner, 1993; Avner et al., 1992).

We discovered that a missense mutation in *furinb* is responsible for the *s741* mutant phenotypes. Furin is a proprotein convertase that functions to cleave latent precursor proteins and convert them to their active state (Denault and Leduc, 1996). *Furin*-null mice die between E10.5-E11.5 due to severe heart malformation and failure of axial rotation (Roebroek et al., 1998). In *s741* mutants, the missense mutation in *furinb* leads to a valine to alanine substitution in the cytosolic domain, which is known to control the trafficking of Furin protein in mammalian cells (Thomas, 2002). Consistently, we found that the mutant Furinb-GFP fusion protein was mislocalized in the cytoplasm of the biliary cells and caused WT cells to cluster. Furin cycles between the trans-golgi network, cell surface, and endosomes to cleave distinct substrates depending on where it is located (Molloy et al., 1999). Known Furin substrates including TGFβ1 (Clotman and Lemaigre, 2006; Dubois et al., 1995), Notch1 receptor (Kitade et al., 2013; Logeat et al., 1998), and IntegrinαV (Lehmann et al., 1996; Patsenker et al., 2008) play important roles in bile duct development. Our transcriptomic analyses did not detect statistically significant changes in Notch, integrin, or TGFβ pathways, likely due to compensation from other proprotein convertases. However, we cannot exclude the possibility that slight perturbations of multiple pathways collectively contribute to the cystic phenotype.

Our study demonstrated the crucial role of ER stress in hepatic cystogenesis. The mislocalized Furinb protein triggered ER stress in *s741* mutant biliary cells. Their cystic phenotype was exacerbated by the ER stress activator Tm and attenuated by the ER stress inhibitor 4-PBA. Increased ER stress can be a general mechanism for hepatic cystogenesis across different vertebrate species. Elevated ER stress has been observed in the liver tissue from PLD patients and PCK rats, as well as in primary cultures of human and rat cystic cholangiocytes (Santos-Laso et al., 2020). Chronic treatment of PCK rats with 4-PBA decreased the volumes of liver cysts. It is noteworthy that most genes identified in isolated PLD (i.e., *PRKCSH*, *SEC63*, *ALG8*, *ALG9, SEC61B*, and *GANAB*) encode proteins that are located in ER and participate in protein biogenesis. It has been speculated that mutations in these genes lead to cystogenesis by affecting posttranslational modulation of PC1 (Besse et al., 2017). Here we propose an alternative hypothesis that mutations in the ER-associated genes may cause a broad perturbation of protein folding, maturation and trafficking, which provokes ER stress. The resulting impairment in cholangiocyte physiology and behaviors results in cyst formation.

We detected an augmentation of mTORC1 pathway in *s741* mutants and showed that chemical inhibition of mTORC1 partially suppressed cyst formation. In patients with ADPKD and ARPKD, the epithelium lining of the hepatic cysts exhibited a markedly increase in mTOR activation compared to control (Becker et al., 2010; Qian et al., 2008). In *Pkd2*-knockout mice, rapamycin decreases liver cystic area and cholangiocyte proliferation through inhibition of insulin-like growth factor signaling and VEGF secretion (Spirli et al., 2010). Clinical trials have been conducted to test the efficacy of mTOR inhibitor treatment in PKD but the results are conflicting (Wuthrich and Mei, 2014). It is thought that high dosage of rapamycin may be needed to reduce cystogenesis while the risk of serious side effects will also increase. Characterizing the mechanisms underlying rapamycin’s action on cystogenesis will discover additional therapeutic targets. Although rapamycin is a potent inducer of autophagy (Noda and Ohsumi, 1998), increasing autophagy alone does not suppress cystogenesis. Treatment with trehalose, which induces autophagy by blocking glucose transport and is independent of mTOR (Mardones et al., 2016), did not reduce the number of hepatic cysts in *s741* mutants. Instead, our data suggest that rapamycin may reduce cyst formation by suppressing ER stress. mTOR signaling has also been previously linked to ER stress. Activation of mTORC1 by deletion of the tuberous sclerosis complex proteins TSC1 or TSC2 causes constitutive ER stress *in vitro* (Ozcan et al., 2008). Liver-specific overexpression of SIRT1, an mTOR inhibitor, attenuated ER stress and insulin resistance in mice (Li et al., 2011). Lastly, as an immunosuppressant, rapamycin may further alleviate biliary injury by decreasing the expression of proinflammatory genes in *s741* mutants.

Our study provides an excellent example of how forward genetic screen elucidates biological phenomena that cannot be predicted from the function of known genes. *furinb* knockout zebrafish and liver-specific knockout of Furin in mice do not cause obvious liver injury (Essalmani et al., 2011; Roebroek et al., 2004), likely due to the redundancy of multiple proprotein convertase family members. The *s741* mutation is a neomorphic allele and only causes cysts in the homozygous mutant or when it was overexpressed in WT by transgene. Given that mutations in the known disease genes only account for 50% of clinical cases of PLD, forward genetic screen in model organisms holds the potential to discover novel genes and mechanisms for hepatic cystogenesis.

## Methods and materials

### Zebrafish

Wild type (WT), furinb^s741+/-^, furinb^ci204+/-^, muc2.2^ci205+/-^, pkd1a^zf1067+/-^, pkd1b^zf1070+/-^ (Zhu et al., 2017), Tg(EPV.Tp1-Mmu.Hbb:EGFP)^um14^/Tg(Tp1:GFP) (Lorent et al., 2010; Parsons et al., 2009), Tg(EPV.Tp1-Mmu.Hbb:hist2h2l-mCherry)^s939^/Tg(Tp1:H2B- mCherry) (Ninov et al., 2012), Tg(mpeg1:YFP)^w200Tg^ (Roca and Ramakrishnan, 2013), and Tg(EPV.Tp1-Mmu.Hbb:mCherry-CAAX)^s733^/Tg(Tp1:ras-mCherry) (Pestel et al., 2016) zebrafish were raised and maintained under standard laboratory conditions (Westerfield, 2007) in accordance with the Guide for the Care and Use of Laboratory Animals (National Institutes of Health publication 86-23, revised 1985) and approved by the Institutional Animal Care and Use Committee at CCHMC. Animals of both genders were studied.

### Whole-genome sequencing and genotyping

Whole-genome sequencing was conducted at the UCSF Genomics Core as described (Leshchiner et al., 2012). Sequencing was performed on pools of 50 *s741* mutants and 50 phenotypically WT siblings based on the morphology of intrahepatic bile ducts revealed by *Tg(Tp1:*GFP) transgene expression. Genomic DNA was prepared using the DNeasy blood and tissue kit (Qiagen, Germantown, MD) and sequenced on Illumina HiSeq 2000 (Illumina, San Diego, CA). Data was analyzed using SNPtrack (http://genetics.bwh.harvard.edu/snptrack/). The *s741* mutation was genotyped by PCR on genomic DNAs using the following primers: forward 5’- TTCTGTCGGAGGACCAAACT -3’, and reverse 5’-ACACACACACACCCACTGGT -3’. The 499 bp PCR product was cut with restriction enzyme Bsp1286I (New England BioLabs, Ipswich, MA). The mutant product was cut into 211 and 288 bp fragments, while the WT product remained uncut at 499 bp.

### *In situ* hybridization and histology

Whole-mount *in situ* hybridization was performed as described (Thisse et al., 1993). To generate a DNA template for making the *furinb* anti-sense RNA *in situ* probe, a fragment of the *furinb* open reading frame was amplified from the total complementary DNA (cDNA) of 96 hpf WT larvae using the following primers: forward 5’- CCGATGACAAGTGACACCAC-3’, and reverse 5’- CACCTCTGTGCTGGAAATGA-3’. Images were obtained on a Carl Zeiss Discovery V8 stereomicroscope using a Zeiss Axiocam MRc5 camera (Carl Zeiss, Oberkochen, Germany). Hematoxylin and eosin (H&E) staining was performed on 6 μm paraffin sections as described previously (Ellis and Yin, 2017), and images were obtained using a Carl Zeiss AxioObserver.Z1 compound microscope. 5 fish per genotype were examined.

### Immunofluorescence

Immunofluorescence staining on whole-mount zebrafish larvae and 150 μm vibratome sections was performed as described (Field et al., 2003; Trinh and Stainier, 2004). 10 fish per genotype were examined for each experiment. Primary and secondary antibodies are listed in Supplemental Table S1. The samples were imaged on a Nikon A1Rsi inverted confocal microscope (Nikon Instruments, Melville, NY) at the Confocal Imaging Core at CCHMC. Imaging processing and quantification were conducted using Imaris software (Bitplane, Concord, MA).

### BODIPY Feeding Assay

At 121 hpf, 20 WT and 20 *s741* mutant larvae were incubated with 6.4 μM BODIPY FL C_5_ (Invitrogen, Waltham, MA) that was suspended in an egg yolk feed at room temperature with agitation for 7 hours. Animals were imaged live on a confocal microscope for assessment of fluorescence in the gallbladder and liver (Carten et al., 2011; Pham and Yin, 2019).

### Transmission Electron Microscopy (TEM)

At 96 hpf, 3 WT and 3 *s741* mutant larvae were fixed in 3% glutaraldehyde in 0.175 M sodium cacodylate buffer (pH7.4) at 4°C overnight, and stained with 1% aqueous uranyl acetate and lead citrate (Pham and Yin, 2019). Thin sections were imaged on a Hitachi H7650 microscope (Hitachi, Ltd., Tokyo, Japan) using the AMT digital camera and AMT image V700 software (Advanced Microscopy Techniques, Corp. Woburn, MA).

### Time-lapse live imaging

At 80 hpf, WT or *s741* mutant larvae expressing *Tg(Tp1:*GFP) were mounted in 1% low-melting agarose in a customized imaging chamber filled with 80 mL egg water containing 0.01% Tricaine (Sigma Aldrich, St. Louis, MO). The chamber was maintained at 28°C for the duration of the time lapses. 40 μm Z-stacks of epifluorescent images of the intrahepatic bile ducts were collected at 8-minute intervals for 14 hours on a Nikon A1R Multiphoton upright confocal microscope. We used a Nikon Apo LWD 25x1.10W DIC N2 objective, an xy resolution of 512X256, and a step size of 1.2 μm. After the recordings, the larvae were genotyped by PCR. Image processing was conducted using Imaris software. 3 WT and 3 mutants were recorded.

### Genetic mosaic analysis by cell transplantation

Cell transplantation was performed as described (Yamashita et al., 2002). The WT embryos expressed *Tg(Tp1:*ras-mCherry) and the *s741* embryos expressed *Tg(Tp1:*GFP). To target the donor cells to the endoderm, 200 pg of *casanova/sox32* mRNA was injected into the donor embryos at the 1-cell stage (Kikuchi et al., 2001; Reiter et al., 2001). Between 4 and 5 hpf, 30-50 donor cells were aspirated from a donor embryo and transplanted into the margin of a host embryo. The donors were genotyped by PCR immediately following transplantation. The host animals were fixed at 120 hpf and the morphology of the host and donor biliary cells was examined by confocal microscopy.

### Chemical treatments

*s741* mutant larvae and their WT and heterozygous siblings expressing *Tg(Tp1:*GFP) were treated with 0.05% DMSO, 5 μM rapamycin (EMD Millipore, Billerica, MA), 10 μM A769662 (Abcam, Waltham, MA), 2% D-(+)-Trehalose dihydrate (Sigma Aldrich), 10 μM 4-Phenylbutyric acid (Sigma Aldrich), or 1mg/mL Tunicamycin (Calbiochem/EMD Chemicals, San Diego, CA) in egg water in the dark from 72 to 120 hpf. They were fixed in 4% PFA and genotyped by PCR.

### Furinb construct design and microinjection

To rescue *s741* phenotypes using WT *furinb* mRNA, a full-length *furinb* cDNA was amplified by PCR from WT total cDNA and inserted into the RNA synthesis plasmid pCS2+. Sense RNA was synthesized from the XhoI-linearized plasmid using the Ambion mMessage mMachine T7 transcription kit (Invitrogen). 20-50 pg of capped *furinb* RNA was injected into the yolk of embryos from *s741* heterozygotes incrosses at the 1-4 cell stage. The injected larvae were fixed in 4% PFA at 96 hpf and genotyped by PCR.

The *Tg(Tp1:wt-furinb-EGFP)* and *Tg(Tp1:s741-furinb-EGFP)* transgenic constructs were generated using the multisite gateway-based Tol2kit (Kwan et al., 2007). The constructs were individually injected into the 10 WT *Tg(Tp1:ras-mCherry)* embryos at the one-cell stage and imaged live using confocal at 120 hpf.

### Statistical analysis

Two-tailed student’s *t*-test and one-way ANOVA with a Tukey’s post-hoc test were performed using GraphPad Prism (GraphPad Software, San Diego, CA).

## Acknowledgement

We would like to thank Dr. Matthew Kofron at CCHMC Confocal Imaging Core for assistance with confocal imaging and post analyses, Drs. Stacey Huppert and Jorge Bezerra for discussions, Drs Adam Miller, Breanne Harty, and Kelly Monk for technical advices on whole genome sequencing, Dr. Zhaoxia Sun for the Arl13b antibody, and Bryan Donnelly for advice on biochemical assays. We also acknowledge CCHMC Veterinary Services for fish care, and Ryan Stefancik, Allison Ross, Zenab Saeed, and Hannah Nartker for research assistance.

