## Supplemental material for "A novel missense mutation in the proprotein convertase gene *furinb* causes hepatic cystogenesis during liver development in zebrafish"

**Supplemental method:**

**Generation of mutant zebrafish strains using CRISPR/Cas9-based genome editing**

To generate mutants with a deletion affecting *muc2.2* similar to what is seen in *s741* mutant, two sgRNAs (CRISPR1: 5’- GGAAGACAATCCCGTTGAAC -3’ and CRISPR2: 5’-GGCAGAAATGGCTGAATATC-3’) flanking the targeted area were manually picked (Supplemental Figure S2B). To make *furinb* loss-of-function mutants, the CRISPR target sequence (5’ – GGGCCGTCATATTGAAGGG – 3’) was identified using the CHOPCHOP website: <http://chopchop.cbu.uib.no/> (Supplemental Figure S3A). Synthesis of sgRNAs and *cas9* mRNA (Chang et al., 2013) was conducted as described (Gagnon et al., 2014). Each sgRNA (300 pg) was co-injected with *cas9* mRNA (500 pg) into WT embryos at the one-cell stage. Genomic DNAs were extracted from a pool of 20 one-day-old embryos and somatic mutations were confirmed by PCR and Sanger sequencing. The injected embryos were raised to adulthood (F0) and screened for germline-transmitting founders (Talbot and Amacher, 2014). The exact mutations were determined by Sanger Sequencing. F0 founders were outcrossed with WT fish to obtain F1 offspring. Heterozygous F1 adults were outcrossed with WT to obtain F2 generation. All experiments involving *mucin2.2^ci205^* and *furinb^ci204^* CRISPR mutants were performed on the animals of F3 or subsequent generations.

To genotype the *muc2.2* deletion mutation, the following PCR primers were used: forward 5’-TGTTCTTGCCTTGTGGTTCA-3’ and reverse 5’-TCGCATCTGATTTCTGGACA-3’. The PCR yielded a 454 bp fragment in WT animals, but no product in the mutants. The *furinb^ci204^* mutation was genotyped by PCR using the primers: forward 5’-CCAGTATTTCTTCCCTGACACTC-3’ and reverse 5’-CAGCATTACATCCACATACACAAG-3’. The 381 bp PCR product was cut with restriction enzyme HpyAV (New England BioLabs, Inc., Ipswich, MA). The WT product was cut into 251 and 128 bp fragments, while the mutant product remained uncut at 381 bp.

**EdU proliferation analysis** and **TUNEL assay**

Proliferating cells were detected using Click-iT EdU Imaging Kit (Thermofisher Scientific, Waltham, MA). Whole-mount TUNEL reaction was conducted using ApopTag Red In Situ Apoptosis Detection Kit according to the manufacturer’s instructions (EMD Millipore, Billerica, MA). Immunofluorescence with anti-GFP antibody was performed after TUNEL staining to detect *Tg(Tp1:*GFP) transgene expression in the biliary cells.

**Reverse transcription PCR for detecting *muc2.2* transcripts in adult and larval zebrafish**

Total mRNAs were prepared using TRIZOL (Sigma Aldrich, St. Louis, MO) from adult WT testes, a pool of ten 4-day-old WT larvae, a pool of ten 4-day-old WT and *s741+/-* larvae, and a pool of ten 4-day-old *s741-/-* mutants. Total mRNAs from a pool of 20 4.5-day-old WT or *s741+/-* larval livers, and a pool of 20 4.5-day-old *s741-/-* larval livers were prepared using PicoPure^TM^ RNA isolation kit (Thermo Fisher Scientific). cDNAs were synthesized by using Superscript III First-Strand Synthesis System (Thermo Fisher Scientific). *muc2.2* PCR was performed using the following primers: 5’- TGATGCTCCAGACAAAACCA-3’ and 5’- AAATTGCTTTGTTCCCTCCA-3’.

**RNA-seq and qPCR**

For transcriptomic analysis, total mRNAs were prepared from pools of 40 dissected livers from *s741* mutants and WT siblings by using PicoPure^TM^ RNA isolation Kit (Thermo Fisher Scientific) and submitted to CCHMC DNA core for library preparation and RNA-sequencing. FASTQ files were analyzed using FASTQC (Andrews, 2010) to ensure uniform read quality (phred>30). Paired end reads were aligned using Star v2.3 (Dobin et al., 2013) to the zebrafish genome GRCz10. The mapped reads were counted using htseq-count (v0.6.0) and gene models from Ensembl transcriptome (Howe et al., 2013). Analysis of differential gene expression was performed using DESeq2 (Love et al., 2014). Principal component analysis was conducted on the regularized log transformed values. Heatmaps were generated using pheatmap package (Kolde, 2012). Orthology to human genes was determined using Ensembl (Collins et al., 2012) and supplemented with orthologs identified by DIOPT (Hu et al., 2011). GSEA was performed using pre-ranked genes based on log2FC (Subramanian et al., 2005).

For qPCR experiments, total mRNAs were prepared from a pool of at least 20 dissected larval livers of each genotype. Total mRNA extraction, cDNA synthesis, and qPCR reactions were performed as described (Zhang et al., 2016). All qPCR reactions were run in triplicate on the 7900HT fast real-time PCR system (Applied Biosystems, Foster City, CA). The relative expression of each gene was determined after normalization to the expression of the housekeeping gene *eef1a1l1* using the relative standard curve method (Larionov et al., 2005). Data analyses were performed using GraphPad Prism Software (GraphPad Software, San Diego, CA). The optimized primers targeting each gene are listed in Supplemental Table S2.

**Supplementary Figures**

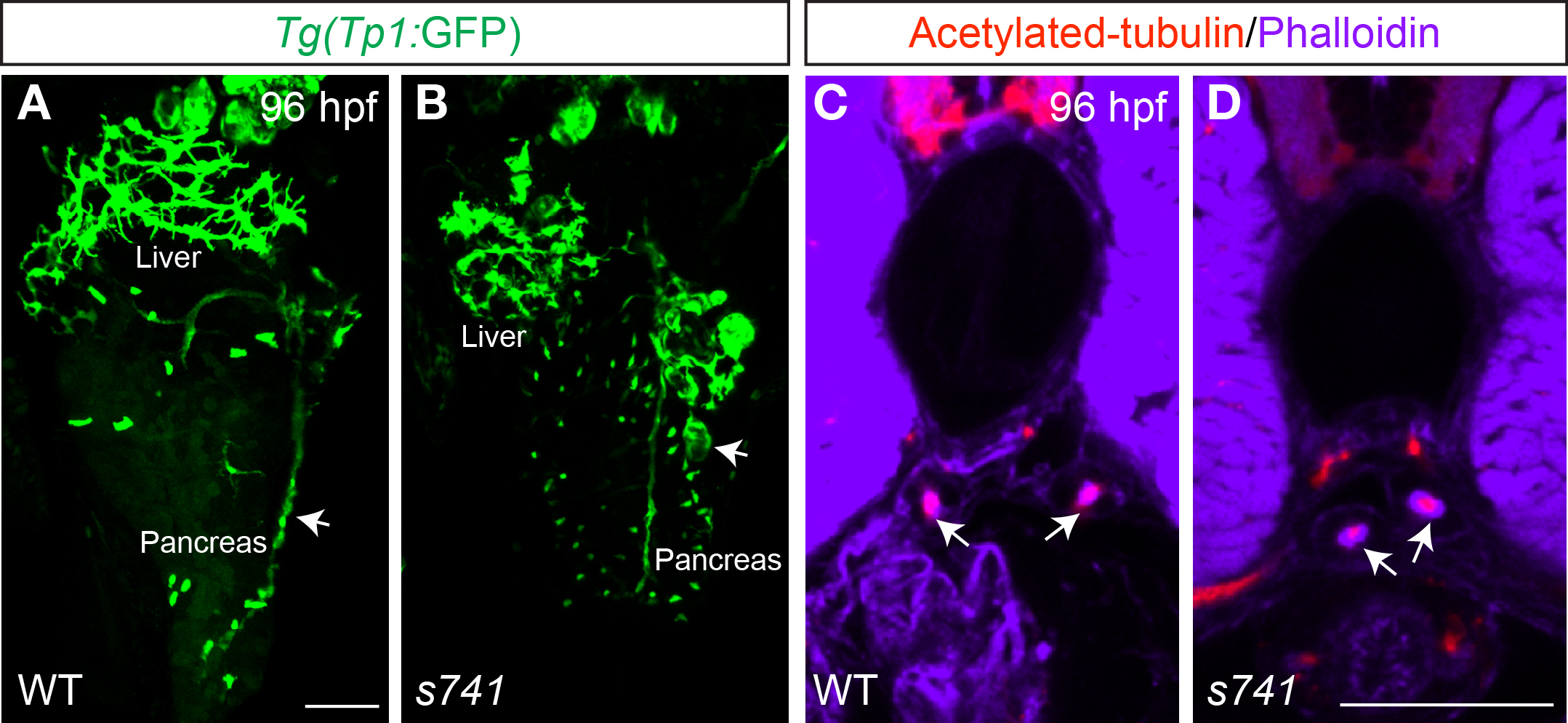

**Supplemental Figure S1.** *s741* mutants develop cysts in the pancreas but not in the pronephric ducts. (A,B) Confocal three-dimensional projections of WT and *s741* mutant zebrafish expressing *Tg(Tp1:*GFP). White arrows point to the pancreatic ducts. *s741* mutants form *Tg(Tp1:*GFP)+ nodules in the pancreas. Ventral views, anterior is to the top. (C,D) Confocal single plane images showing vibratome sections of WT and *s741* mutant zebrafish stained with phalloidin (purple) for F-actin and acetylated-tubulin antibody (red) for stabilized microtubules. White arrows mark the pronephric ducts. The mutant pronephric ducts do not seem to contain cysts. Serial sections through the entire length of the pronephric ducts were examined in five WT and five mutant larvae. Transverse sections, dorsal is to the top. hpf, hours post fertilization. Scale bars, (A,B) 70 μm; (C,D) 50 μm.

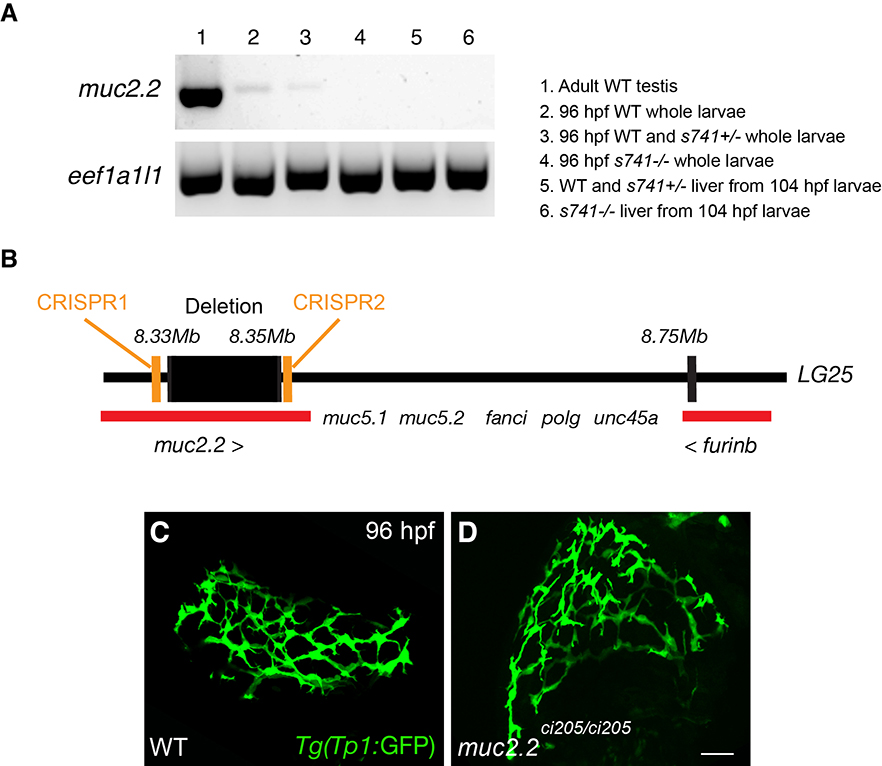

**Supplemental Figure S2.** The *muc2.2* deletion mutation unlikely contributes to the formation of hepatic cysts in *s741* mutants. (A) Reverse transcription PCR to detect *muc2.2* transcript expression in the cDNAs isolated from adult WT male testes, pooled WT, *s741* heterozygous, and homozygous mutant larvae at 96 hpf, and pooled livers from WT, *s741* heterozygous, and homozygous mutant larvae at 96 hpf. The housekeeping gene *eef1a1l1* was used as the loading control. (B) Diagram of the region on Linkage Group (LG) 25 that contains the *muc2.2* and *furinb* genes. Their coding regions are highlighted in red. The *muc2.2* deletion found in *s741* mutants is marked with the black box. The two CRISPR target sites are marked in orange. (C,D) Confocal three-dimensional projections of WT and *muc2.2^ci205^* CRISPR mutant larvae at 96 hpf. *Tg(Tp1:*GFP) expression labels the intrahepatic bile ducts. Ventral views, anterior is to the top. Scale bar, 30 μm.

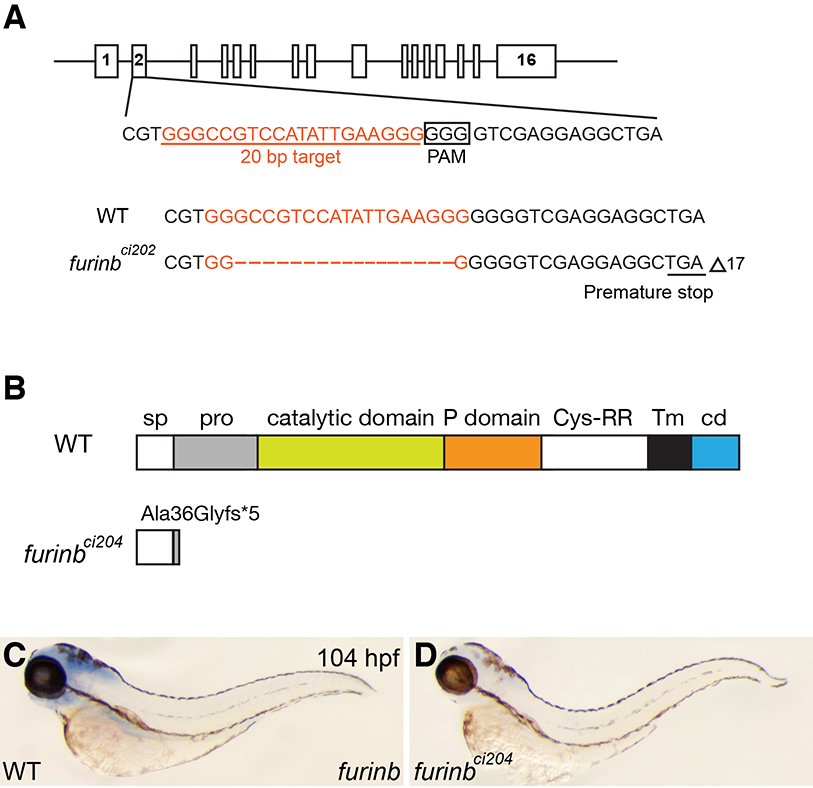

**Supplemental Figure S3.** Generation of *furinb^ci204^* mutants by CRISPR/Cas9 genome-editing technology. (A) Schematic representation of the zebrafish *furinb* locus along with the CRISPR target site, PAM motif, and WT and indel sequences. (B) Putative domain diagram of Furinb protein in WT and *furinb^ci204^* mutant. The predicted changes in CRISPR mutant protein are shown. Sp, signal peptide; pro, pro-domain; Cys-RR, cysteine-rich region; Tm, transmembrane domain; cd, cytoplasmic domain. (C,D) Whole-mount *in situ* hybridization detected an evident reduction of *furinb* mRNA expression in *furinb^ci204^* mutant (D) compared to WT (C). NBT/BCIP staining was stopped immediately after the signal became apparent in some larvae. The larvae with positive signal (20/25) and those without (5/20) were genotyped by PCR. The positive larvae were either WT or *furinb^ci204+/-^* heterozygotes and the negative larvae were all *furinb^ci205-/-^* homozygotes.

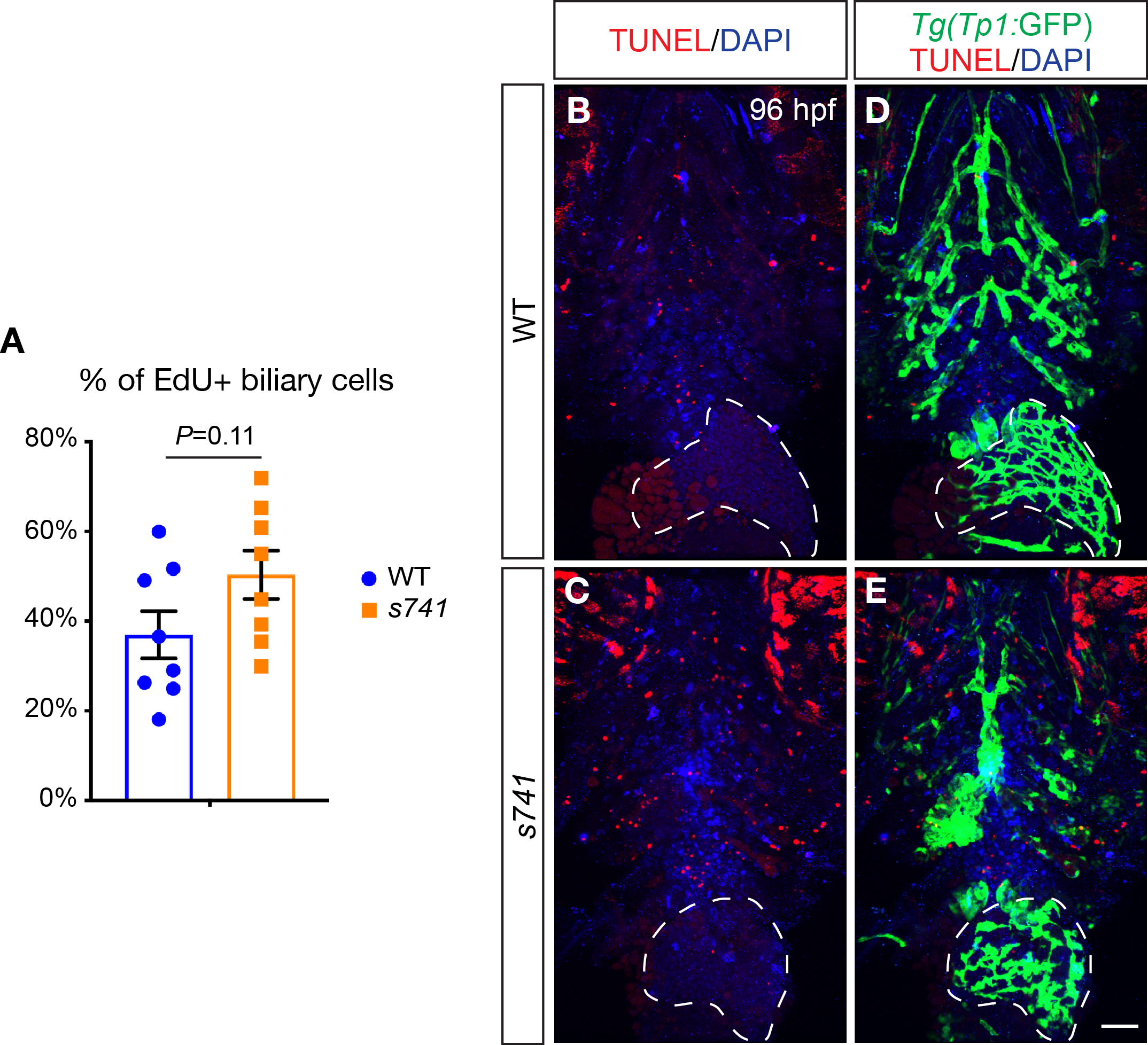

**Supplemental Figure S4.** Dysregulation of biliary cell proliferation or apoptosis does not cause hepatic cyst formation in *s741* mutants. (A) Percentages (mean±s.e.m.) of *Tg(Tp1:*GFP)+ intrahepatic biliary cells that incorporated proliferation marker EdU between 80 and 120 hpf. Statistical significance was calculated by two-tailed student’s *t*-test. Each dot represents an individual liver. (B-E) Confocal three-dimensional projections of WT and *s741* mutant zebrafish stained by TUNEL assay. (B,C) show TUNEL labeling (red) that marked apoptotic cells and DAPI (blue) that stained nuclei. (D,E) are the same fish as shown in (B,C) but with *Tg(Tp1:*GFP) expression (green) labeling the intrahepatic biliary cells. In both WT and mutants, substantial numbers of TUNEL-positive cells can be seen in the pharyngeal arches; however, there were very few apoptotic cells in the livers (outlined by the dashed lines). Five WT and five mutant fish were examined. Ventral views, anterior is to the top. Scale bars, (B-E) 50 μm.

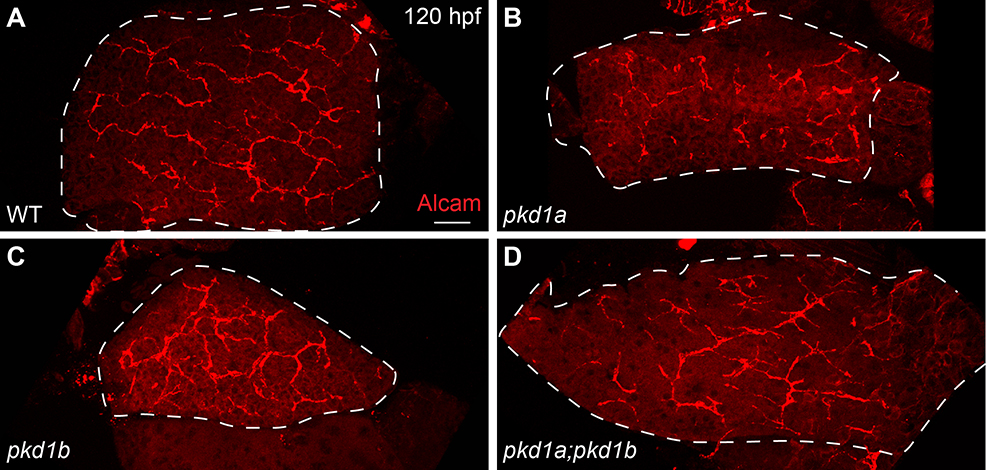

**Supplemental Figure S5.** The intrahepatic biliary cells in zebrafish *pkd1* mutant larvae do not form cysts. (A-D) Confocal three-dimensional projections of livers in WT (A), *pkd1a* (B) and *pkd1b* (C) individual mutants, and *pkd1a;pkd1b* compound homozygous mutants (D). The biliary marker Alcam showed that none of the mutants had biliary nodules as seen in *s741* mutants at 120 hpf. 10 fish per genotype were examined. Ventral views, anterior is to the top. Scale bar, 30 μm.

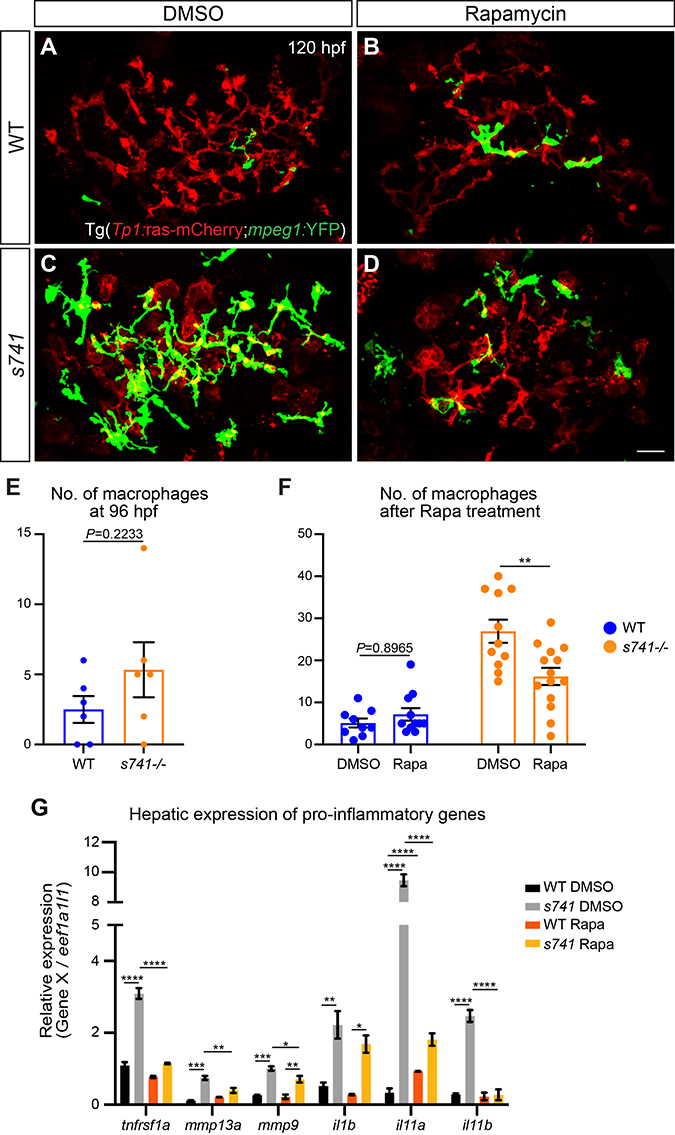

**Supplemental Figure S6.** Rapamycin treatment decreases inflammatory responses in *s741* mutants. (A-D) Confocal three-dimensional projections of WT and *s741* mutant livers after DMSO or 5 μM rapamycin treatment from 72 to 120 hpf. The intrahepatic biliary cells are marked by *Tg(Tp1:*ras-mCherry) expression. Macrophages are labeled by *Tg(mpeg1:*YFP) expression. Ventral views, anterior is to the top. Scale bar, 50 μm. (E) Number of macrophages (mean±s.e.m.) in WT and *s741* mutant livers at 96 hpf. (F) Number of macrophages (mean±s.e.m.) in WT and *s741* mutant livers at 120 hpf right after DMSO or rapamycin treatment. In (E,F), each point represents an individual liver. (G) qPCR analyses showing the hepatic expression of pro-inflammatory genes in WT and *s741* mutants after DMSO or rapamycin treatment from 72 to 120 hpf. Triplicates were performed. The results are represented as relative expression levels that are normalized to the housekeeping gene *eef1a1l1* (mean±s.e.m.). Statistical significance in (E,F,G) was calculated by one-way ANOVA and Tukey’s post-hoc test. *, p<0.05; **, p<0.01; ***, p<0.001; ****, p<0.0001. Rapa, rapamycin.

Howe, K., Clark, M.D., Torroja, C.F., Torrance, J., Berthelot, C., Muffato, M., Collins, J.E., Humphray, S., McLaren, K., Matthews, L., McLaren, S., Sealy, I., Caccamo, M., Churcher, C., Scott, C., Barrett, J.C., Koch, R., Rauch, G.J., White, S., Chow, W., Kilian, B., Quintais, L.T., Guerra-Assuncao, J.A., Zhou, Y., Gu, Y., Yen, J., Vogel, J.H., Eyre, T., Redmond, S., Banerjee, R., Chi, J., Fu, B., Langley, E., Maguire, S.F., Laird, G.K., Lloyd, D., Kenyon, E., Donaldson, S., Sehra, H., Almeida-King, J., Loveland, J., Trevanion, S., Jones, M., Quail, M., Willey, D., Hunt, A., Burton, J., Sims, S., McLay, K., Plumb, B., Davis, J., Clee, C., Oliver, K., Clark, R., Riddle, C., Elliot, D., Threadgold, G., Harden, G., Ware, D., Begum, S., Mortimore, B., Kerry, G., Heath, P., Phillimore, B., Tracey, A., Corby, N., Dunn, M., Johnson, C., Wood, J., Clark, S., Pelan, S., Griffiths, G., Smith, M., Glithero, R., Howden, P., Barker, N., Lloyd, C., Stevens, C., Harley, J., Holt, K., Panagiotidis, G., Lovell, J., Beasley, H., Henderson, C., Gordon, D., Auger, K., Wright, D., Collins, J., Raisen, C., Dyer, L., Leung, K., Robertson, L., Ambridge, K., Leongamornlert, D., McGuire, S., Gilderthorp, R., Griffiths, C., Manthravadi, D., Nichol, S., Barker, G., Whitehead, S., Kay, M., Brown, J., Murnane, C., Gray, E., Humphries, M., Sycamore, N., Barker, D., Saunders, D., Wallis, J., Babbage, A., Hammond, S., Mashreghi-Mohammadi, M., Barr, L., Martin, S., Wray, P., Ellington, A., Matthews, N., Ellwood, M., Woodmansey, R., Clark, G., Cooper, J., Tromans, A., Grafham, D., Skuce, C., Pandian, R., Andrews, R., Harrison, E., Kimberley, A., Garnett, J., Fosker, N., Hall, R., Garner, P., Kelly, D., Bird, C., Palmer, S., Gehring, I., Berger, A., Dooley, C.M., Ersan-Urun, Z., Eser, C., Geiger, H., Geisler, M., Karotki, L., Kirn, A., Konantz, J., Konantz, M., Oberlander, M., Rudolph-Geiger, S., Teucke, M., Lanz, C., Raddatz, G., Osoegawa, K., Zhu, B., Rapp, A., Widaa, S., Langford, C., Yang, F., Schuster, S.C., Carter, N.P., Harrow, J., Ning, Z., Herrero, J., Searle, S.M., Enright, A., Geisler, R., Plasterk, R.H., Lee, C., Westerfield, M., de Jong, P.J., Zon, L.I., Postlethwait, J.H., Nusslein-Volhard, C., Hubbard, T.J., Roest Crollius, H., Rogers, J., Stemple, D.L., 2013. The zebrafish reference genome sequence and its relationship to the human genome. Nature 496, 498-503.

Hu, Y., Flockhart, I., Vinayagam, A., Bergwitz, C., Berger, B., Perrimon, N., Mohr, S.E., 2011. An integrative approach to ortholog prediction for disease-focused and other functional studies. BMC Bioinformatics 12, 357.

Kolde, R., 2012. Pheatmap: Pretty heatmaps.

Larionov, A., Krause, A., Miller, W., 2005. A standard curve based method for relative real time PCR data processing. BMC Bioinformatics 6, 62.

Love, M.I., Huber, W., Anders, S., 2014. Moderated estimation of fold change and dispersion for RNA-seq data with DESeq2. Genome Biol 15, 550.

Subramanian, A., Tamayo, P., Mootha, V.K., Mukherjee, S., Ebert, B.L., Gillette, M.A., Paulovich, A., Pomeroy, S.L., Golub, T.R., Lander, E.S., Mesirov, J.P., 2005. Gene set enrichment analysis: a knowledge-based approach for interpreting genome-wide expression profiles. Proc Natl Acad Sci U S A 102, 15545-15550.

Talbot, J.C., Amacher, S.L., 2014. A streamlined CRISPR pipeline to reliably generate zebrafish frameshifting alleles. Zebrafish 11, 583-585.

Zhang, C., Ellis, J.L., Yin, C., 2016. Inhibition of vascular endothelial growth factor signaling facilitates liver repair from acute ethanol-induced injury in zebrafish. Dis Model Mech 9, 1383-1396.

**Table S1 Primary and secondary antibodies used in the study.**

| **Antigen** | **Company** | **Catalog number** | **Dilution** |
| --- | --- | --- | --- |
| Annexa4 (2F11) | Abcam | ab71286 | 1:100 |
| Alcam | ZIRC | zn8 | 1:20 |
| GFP | Aves Lab | GFP-1020 | 1:1000 |
| Prox1 | Millipore | AB5475 | 1:1000 |
| BSEP | Kamiya Biomedical | PC-064 | 1:1000 |
| γ-tubulin | Sigma-Aldrich | T6557 | 1:500 |
| Acetylated-tubulin | Sigma-Aldrich | T6793 | 1:200 |
| Arl13b | Gift from Zhaoxia Sun | NA | 1:200 |
| DAPI | Thermo Fisher Scientific | D1306 | 1:1000 |
| Goat anti-rabbit IgG (H+L), Alexa Fluor 568 | Thermo Fisher Scientific | A11011 | 1:200 |
| Donkey anti-mouse IgG (H+L), Alexa Fluor 647 | Thermo Fisher Scientific | A315571 | 1:200 |
| Goat anti-mouse IgG (H+L), Alexa Fluor 568 | Thermo Fisher Scientific | A11004 | 1:200 |
| Goat anti-chicken IgG (H+L), Alexa Fluor 488 | Thermo Fisher Scientific | A11039 | 1:200 |
| Alexa Fluor 647 phalloidin | Thermo Fisher Scientific | A22287 | 1:100 |
| Rhodamine Phalloidin | Thermo Fisher Scientific | R415 | 1:100 |

**Supplementary Table S2.** qPCR primers

| Gene | Accession Number | Primers |
| --- | --- | --- |
| *furinb* | NM_001045109 | 5’- ACGCTCTCCATCAGCAGCAC -3’  5’- GTTTCTCGTTGAGGTTTCCGC -3’ |
| *atf4a* | NM_213233 | 5’- TTAGCGATTGCTCCGATAGC-3’  5’- GCTGCGGTTTTATTCTGCTC-3’ |
| *atf6* | NM_001110519 | 5’- CTGTGGTGAAACCTCCACCT-3’  5’- CATGGTGACCACAGGAGATG-3’ |
| *ddit3/chop* | NM_001082825 | 5’- AAGGAAAGTGCAGGAGCTGA-3’  5’- TCACGCTCTCCACAAGAAGA-3’ |
| *dnajc3a* | NM_199610 | 5’- TCCCATGGATCCTGAGAGTC-3’  5’- CTCCTGTGTGTGAGGGGTCT-3’ |
| *edem1* | NM_201189 | 5’- ATCCAAAGAAGATCGCATGG-3’  5’- TCTCTCCCTGAAACGCTGAT-3’ |
| *eef1a1l1/ef1a* | NM_131263 | 5’-CTTCTCAGGCTGACTGTGC-3’  5’-CCGCTAGCATTACCCTCC-3’ |
| *hsp90b1/grp94* | NM_198210 | 5’- AGCAAGACCGAGACCGTAGA-3’  5’- CTCCCAATCCCACACAGTCT-3’ |
| *hspa5/bip* | NM_213058 | 5’- AAGAGGCCGAAGAGAAGGAC-3’  5’- AGCAGCAGAGCCTCGAAATA-3’ |
| *il1b* | NM_212844 | 5’-TGGACTTCGCAGCACAAAATG-3’  5’-GTTCACTTCACGCTCTTGGATG-3’ |
| *il11a* | XM_693882 | 5’-CTCCTTCTGACGCTGACTCG-3’  5’-TTGCGAAGTCACTGGCTCTG-3’ |
| *il11b* | XM_021468285 | 5’-GCTATCATCCCTGCCCTCAC-3’  5’-GATCTCGGGTGCTGTCTGTC-3’ |
| *mmp9* | NM_213123 | 5’-GAAGCGTTACGGCTACGT-3’  5’-TTCCATGTCTGGCGAATA-3’ |
| *mmp13a* | NM_001290479 | 5’-GTGATGAAAAAGCCCCGCTG-3’  5’-CATCGTCGAAATGAGCGTCG-3’ |
| *tnfrsf1a* | NM_213190 | 5’-AGCATTCCCCCAGTCTTTTT-3’  5’-GCAGGTGACGATGACTGAGA-3’ |
| *xbp1-s* | NM_131874 | 5’- TGTTGCGAGACAAGACGA-3’  5’- CCTGCACCTGCTGCGGACT-3’ |
| *xbp1-t* | NM_131874 | 5’- GGGTTGGATACCTTGGAAA-3’  5’- AGGGCCAGGGCTGTGAGTA-3’ |
